# EpiGePT: a Pretrained Transformer model for epigenomics

**DOI:** 10.1101/2023.07.15.549134

**Authors:** Zijing Gao, Qiao Liu, Wanwen Zeng, Rui Jiang, Wing Hung Wong

**Author notes:** The first two authors contributed equally.

## Abstract

The inherent similarities between natural language and biological sequences have given rise to great interest in adapting the transformer-based large language models (LLMs) underlying recent breakthroughs in natural language processing (references), for applications in genomics. However, current LLMs for genomics suffer from several limitations such as the inability to include chromatin interactions in the training data, and the inability to make prediction in new cellular contexts not represented in the training data. To mitigate these problems, we propose EpiGePT, a transformer-based pretrained language model for predicting context-specific epigenomic signals and chromatin contacts. By taking the context-specific activities of transcription factors (TFs) and 3D genome interactions into consideration, EpiGePT offers wider applicability and deeper biological insights than models trained on DNA sequence only. In a series of experiments, EpiGePT demonstrates superior performance in a diverse set of epigenomic signals prediction tasks when compared to existing methods. In particular, our model enables cross-cell-type prediction of long-range interactions and offers insight on the functional impact of genetic variants under different cellular contexts. These new capabilities will enhance the usefulness of LLM in the study of gene regulatory mechanisms. We provide free online prediction service of EpiGePT through http://health.tsinghua.edu.cn/epigept/.

## Introduction

A fundamental but largely unresolved problem in genomics is to decode the information residing in the non-coding part of the human genome^1^. It remains incompletely understood how regulatory elements govern gene expression in different contexts^1^, and how noncoding variants may disrupt the underlying regulatory syntax of DNA^2^. Fortunately, recent advances in epigenome sequencing^3, 4^ have resulted in the accumulation of data useful for the study of these questions, including chromatin accessibility, DNA methylation, histone modifications, and 3D chromatin interaction. Thus, there is great interest in performing systematic analysis of these data to enhance our ability to interpret the non-coding part of the genome^5–11^.

The inherent similarities between natural language and biological sequences has also stimulated interest in developing large language models (LLM) for the interpretation of genome sequences^12^. As is well known, the development of large language model (LLM) has been the main driving force behind many recent breakthroughs in artificial intelligence such as ChatGPT. The architecture of the LLM is typically a multilayer transformer network, and the model is trained on a very large corpus of natural language data. Such pre-trained models can be readily tailored or adapted to various downstream tasks. Considering DNA sequences as the texts in the genomic language, similar transformer-based approaches have been used to model DNA sequences^13, 14^. For example, the Enformer model^15^ takes the DNA sequence of a large genomic region as input and predict thousands of epigenomic features across cellular contexts covered by the training data. Although already useful in many applications, such models relying on only DNA sequences as input are not capable of predicting the function of sequences in new cellular contexts. Furthermore, despite the importance of 3D chromatin contacts in gene regulation, 3D interaction data have not been included in the training of current genomic LLMs. Therefore, there is an urgent need to further develop the core technologies of genomic LLMs to overcome these limitations.

In this paper, we present EpiGePT, a transformer-based model for epigenomics prediction with the following new capabilities. First, the inability to make predictions in novel contexts has greatly limited the applicability of current methods, EpiGePT removes this limitation by making both the input and output context-dependent, where the context is represented by a TF-profile vector specifying the expression of key TFs in that context. This choice is motivated by the fact that reference gene expression data are available for many cellular contexts that are important in development and diseases, but for which few epigenomic features have been measured. We note that the reference TF expression profile has been used to represent cellular context in earlier works on accessibility prediction^6, 16^, but this idea has not been explored for the development of genomic LLMs. Second, a new learning algorithm is developed to enable the inclusion of 3D chromatin contact data in the training data. In this way, EpiGePT can predict 3D genome features such as enhancer-promoter interactions that are known to be important for gene regulation but are not modeled in current genomic LLMs. By using a masked training strategy, EpiGePT can be trained on a diverse set of contexts even if different sets of epigenomics signals are available in different contexts. There is a profound difference in training strategy between EpiGePT and current genomic LLMs. Each input genomic region provides an example for training in current LLMs such as the Enformer. In contrast, each combination of input region and cellular context provides an example for training in EpiGePT, thus providing a much larger number of examples available for model training. As for training data sets, since most cellular contexts that have epigenomic data will also have expression data, we can use most available epigenomic data, such as those used by the Enformer, to train our model.

In a series of experiments, we illustrate that our model is superior to existing methods in epigenomic signals prediction, long-range chromatin interaction prediction, as well as the variant effect prediction.

## Results

### Overview of EpiGePT

EpiGePT is a genomic language model for cross-cell-type prediction of chromatin states by multi-task learning based on genome-wide pre-training on epigenomic data (Fig. 1 and Fig. S2). The model is composed of four modules, including a sequence module, a TF module, a transformer module, and a prediction module. The sequence module is responsible for processing the long DNA sequence of interest (e.g., 128 kb) by employing a series of convolutional and pooling blocks (e.g., 5) to extract a comprehensive set of sequence features. The TF module is specifically designed to represent a cellular context by a TF-profile vector, which specifies the state of a few hundred TFs in that context. The features computed by the sequence and TF modules are then fed as input tokens to the transformer module, where each token corresponds to a genomic bin (e.g., a 128 bp window) in the original DNA sequence. The transformer module leverages self-attention mechanisms to learn the relationships among the input bins, enabling the model to make predictions of multiple chromatin states given the context information from the TF module. Importantly, by including a novel loss term that involves the self-attention weights, EpiGePT is capable of learning from data on context-specific chromatin interactions. Since 3D interaction is known to be a key mechanism in gene regulation, the ability to learn from interaction data is an attractive feature of our approach. Finally, the fourth module in EpiGePT is a predictive module which predicts epigenomic signals and chromatin interactions based on the output of the transformer module.

**Fig. 1.**
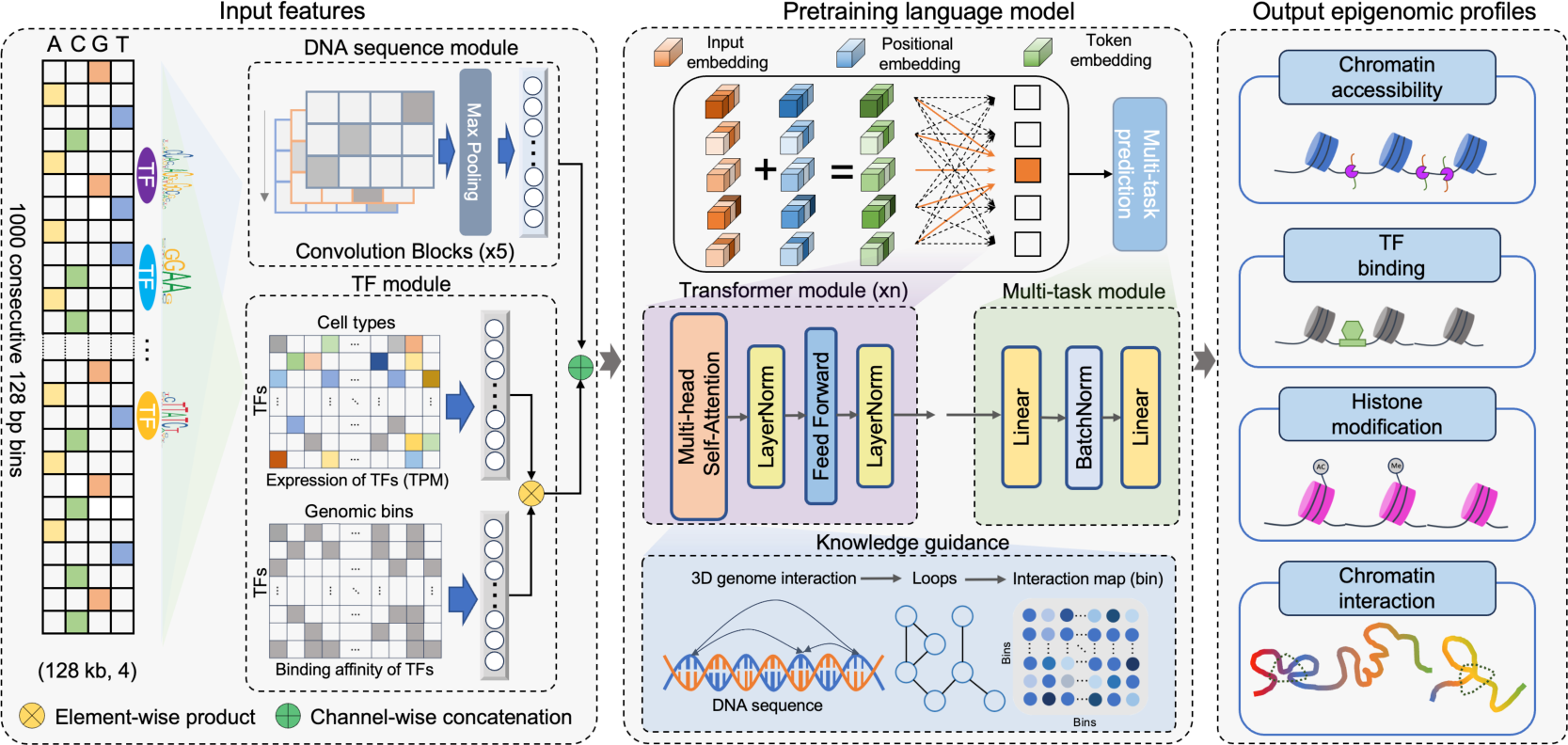
Overview of the EpiGePT model for multiple epigenomic signals prediction. The EpiGePT model consists of four modules, namely the Sequence module, the TF module, the Transformer module, and the Multi-task prediction module. The sequence module comprises multiple layers of convolution applied to the one-hot encoded DNA sequence input. The input sequence length consists of 1000 genomic bins of 128bp for the prediction of multiple signals and 50 bins of 200bp for the prediction of DNase signal alone. The TF module encompasses the binding status and expression of 711 transcription factors. The Transformer module consists of a series of consecutive transformer encoders, while the multi-task module is composed of a fully connected layer. Additionally, the EpiGePT framework integrates an optional knowledge guidance module that enhances the interpretability of the model by incorporating three-dimensional chromatin interaction data into the attention layer, thus improving its understanding of regulatory mechanisms.

### Genome-wide prediction of epigenomic signals

To assess the performance on predicting epigenomic signals, we first compared EpiGePT to task-specific models that are specifically designed for predicting a single epigenomic signal. Taking the chromatin accessibility for instance, the performance of EpiGePT was compared against existing task-specific models such as BIRD^17^, ChromDragoNN^6^, and DeepCAGE^16^. The widely available public DNase-seq^18^ data across 129 cellular contexts on 1,175,374 genomic regions were collected and preprocessed from ENCODE database^19^ (see Methods). Performance is evaluated in three prediction settings: i) “cross-region” setting where the predictive model is tested on new genomic regions not seen in training, ii) “cross-cell type” setting where the model is tested on new cell types, and iii) “cross-both” setting where testing is done on new regions in new cell types (Fig. S1, Supplementary Text S1). In each setting, we employed three evaluation metrics, namely Pearson correlation coefficient (PCC), Spearman correlation coefficient (SCC) and prediction square error (PSE), to assess the similarity between the predicted and true values of the DNase signals (See Methods). The results, presented in Fig. 2a and Fig. S3, showed that EpiGePT consistently outperformed baseline methods including BIRD^17^, and ChromDragoNN^6^ by a relatively large margin under the above settings. For example, EpiGePT achieved a cross-cell type prediction PCC of 0.787, demonstrated a 6.9% higher performance than the best baseline method, ChromDragoNN. In addition, we also evaluated the prediction of binary chromatin accessibility status i.e. predicting whether a peak exists within the corresponding genomic bin (>50% overlap). For binary prediction, EpiGePT again achieved a superior performance with an average auPRC (area under the precision-recall curve) of 0.767 compared to 0.623 of DeepCAGE^16^ and 0.476 of ChromDragoNN^6^ (Fig. 2c). Finally, we compared EpiGePT and ChromDragoNN^6^ in the binary classification of functional regions versus nonfunctional regions, using the functional chromatin status derived from ChromHMM^20^ annotations as ground truth (Supplementary Text S6). EpiGePT achieved an average 8.1% higher auROC (area under the receiver operating characteristic curve) than ChromDragoNN^6^, and an average 2.3% higher macro-auROC than ChromDragoNN^6^ (*p*-value < 0.001 under one-sided Wilcoxon signed rank test) in a finer-grained classification for different types of regulatory elements (Fig. S4). These results demonstrate that EpiGePT provides better predictions than task-specific models.

**Fig. 2.**
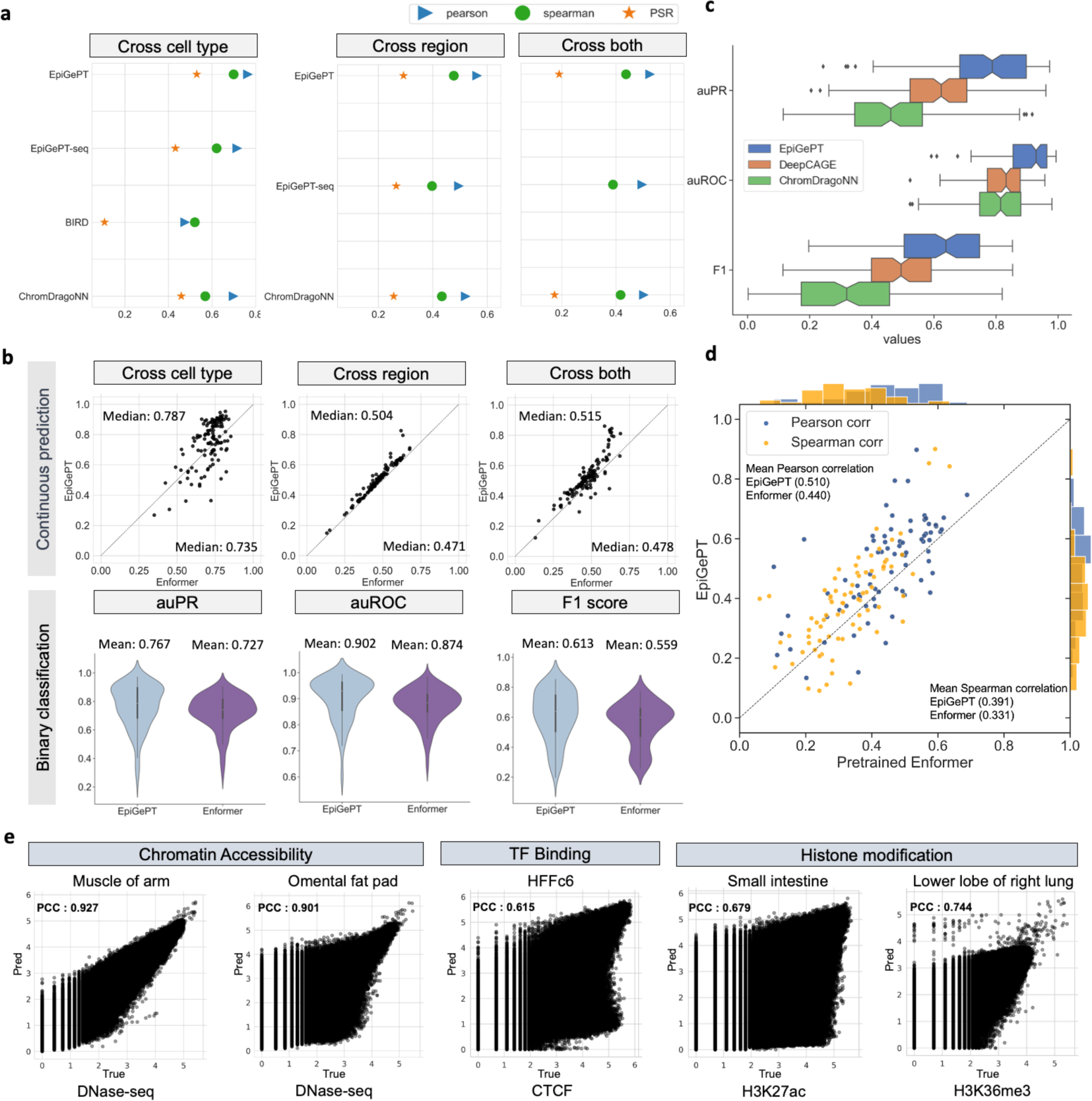
Performance of EpiGePT and baseline methods on the benchmark experiment. **a**, EpiGePT and baseline methods were compared in terms of their regression performance for DNase signal regression across cell types, genomic regions, and combined cell type and genomic regions. **b**, Comparison of EpiGePT and Enformer performance. Each point in the scatter plot represents the performance of Enformer on the data of a specific cell type (x-axis) compared to the performance of EpiGePT (y-axis). The top three graphs represent the prediction of continuous DNase signals (pearson correlation coefficient), while the bottom three graphs represent the binary classification of chromatin accessibility regions. **c**, EpiGePT and baseline methods’ performance on binary prediction of DNase-seq signals. **d**, EpiGePT demonstrates more excellent performance in predicting diverse epigenetic signals across various cell types, compared with the pre-trained Enformer on 78 genomic tracks across 19 unseen cell types. The orange points represent Spearman correlation coefficient, and the blue points represent pearson correlation coefficient. **e**, EpiGePT cross-cell-type predictions compared to experimental signals visualized for a representative example. The predictions specific to DNase are based on the hg19 reference genome, while predictions for multiple epigenomic profiles are conducted using the hg38 reference genome.

Next, we compared EpiGePT with a state-of-the-art genomic LLM, Enformer^15^, in two different ways. First, we trained an Enformer model from scratch with only the aforementioned DNase-seq data (Supplementary Text S5). EpiGePT demonstrates a 3.3% to 5.2% higher performance than Enformer in terms of the median Pearson correlation coefficient under the three prediction settings (Fig. 2b). Second, we compared EpiGePT directly to the pretrained Enformer model provided by the original paper. To do this, we collected eight different epigenomic signals from 104 different cellular contexts (Supplementary Table S4, S6 and S9). We first left out 13 of these contexts where HiChIP data are also available for downstream chromatin interactions validation. Then, EpiGePT model was trained across 72 training cellular contexts (without using HiChIP-based chromatin contacts data in the training) and subsequently compared against pre-trained Enformer on the remaining 19 test cellular contexts, on 15,870 training genomic regions with 128kbp length. Since most of the cellular contexts have missing epigenomic signals, we designed a masked training strategy to handle this issue (See Methods). Under the test cellular contexts, EpiGePT exhibited superior performance with higher PCC than Enformer in 60 out of 78 matched epigenomic signals across 19 test cellular contexts by achieving an average PCC of 0.510, compared to 0.440 of Enformer (Fig. 2d and Fig. S6b). For DNase-seq specifically, the average PCC of EpiGePT reached 0.710 and the average SCC reached 0.664 across 7 cell types, compared to the average PCC of 0.455 and the average SCC of 0.488 of Enformer. In the above comparison, we are in fact comparing out-sample prediction by EpiGePT to in-sample prediction by Enformer. The favorable results achieved by EpiGePT in this experimental setting suggests that our model enables prediction in novel contexts without sacrificing performance. To illustrate the prediction performance further, several tracks of predicted chromatin states and the corresponding ground truth chromatin states were displayed in Fig.2e.

### EpiGePT enables the prediction of chromatin interactions

We examined the capacity of EpiGePT for predicting long-range chromatin interactions, which is important for understanding chromatin architecture and relations between regulatory elements and target genes. We employed several experimental settings to examine the ability of EpiGePT in capturing long-range chromatin interactions. In setting (A), we directly utilized the self-attention weights extracted from the pretrained EpiGePT model (without including HiChIP data in the training) to predict enhancer-promoter (E-P) interactions and silencer-promoter (S-P) interactions. In setting (B), we integrated HiChIP-derived 3D chromatin contacts into the training of the model and then use the model to predict E-P interactions in novel contexts not seen in the training. In setting (C), we designed a pretrain-finetune strategy for EpiGePT model to predict E-P interactions. The results under each setting are discussed below.

Setting (A): prediction by EpiGePT not trained with 3D contact data. In this setting, we use the cell-type specific self-attention scores to predict chromatin interactions, including E-P and S-P interactions (see Methods). Two sets of interactions containing 664 and 5,091 candidate element-gene interactions, obtained by CRISPRi^21^ experiments on K562 cell line, were collected and further filtered and divided into positive and negative samples, for use as ground truths to evaluate E-P prediction performance. In the Gasperini et al^22^. dataset, EpiGePT consistently outperformed Enformer by achieving the highest auPRC in most cases (Fig. 3a). For instance, EpiGePT achieved auPRC of 0.647 to 0.887 for identifying enhancer-gene transcription start site (TSS) pairs in different distance groups (Fig. 3a and Fig. S7). In the Fulco et al.^23^ dataset, EpiGePT also outperformed other competing methods. For example, EpiGePT achieves an auPRC of 0.504, compared to 0.307 of Enformer in the 30-45kbp group (Fig. 3a). Next, to assess performance on S-P interactions., we downloaded putative silencers from the SilencerDB^24^ and used the TSS of annotated nearest gene as the potential target. We selected the same number of negative pairs randomly while conserving the distance distribution. The results show that EpiGePT achieved a better performance in distinguishing positive S-P pairs from negative pairs than Enformer. For instance, EpiGePT achieves an auROC of 0.575 in long-range S-P interactions (32-64kbp) compared to 0.483 of Enformer (Fig. 3b). Finally, to assess performance in predicting chromatin interactions, we collected HiChIP^25^ loops on K562 and GM12878 cell lines from the HiChIPdb^26^. EpiGePT achieves a superior performance by discerning HiChIP loops from randomly selected loops with the same distance distribution. For instance, EpiGePT achieves an auPRC of 0.520 for long range loops (40-64kbp) prediction in GM12878 cell line, surpassing that of Enformer (0.484) by a large margin (Fig. 3g). These results clearly demonstrated the utility of EpiGePT attention scores in capturing functional chromatin interactions.

**Fig. 3.**
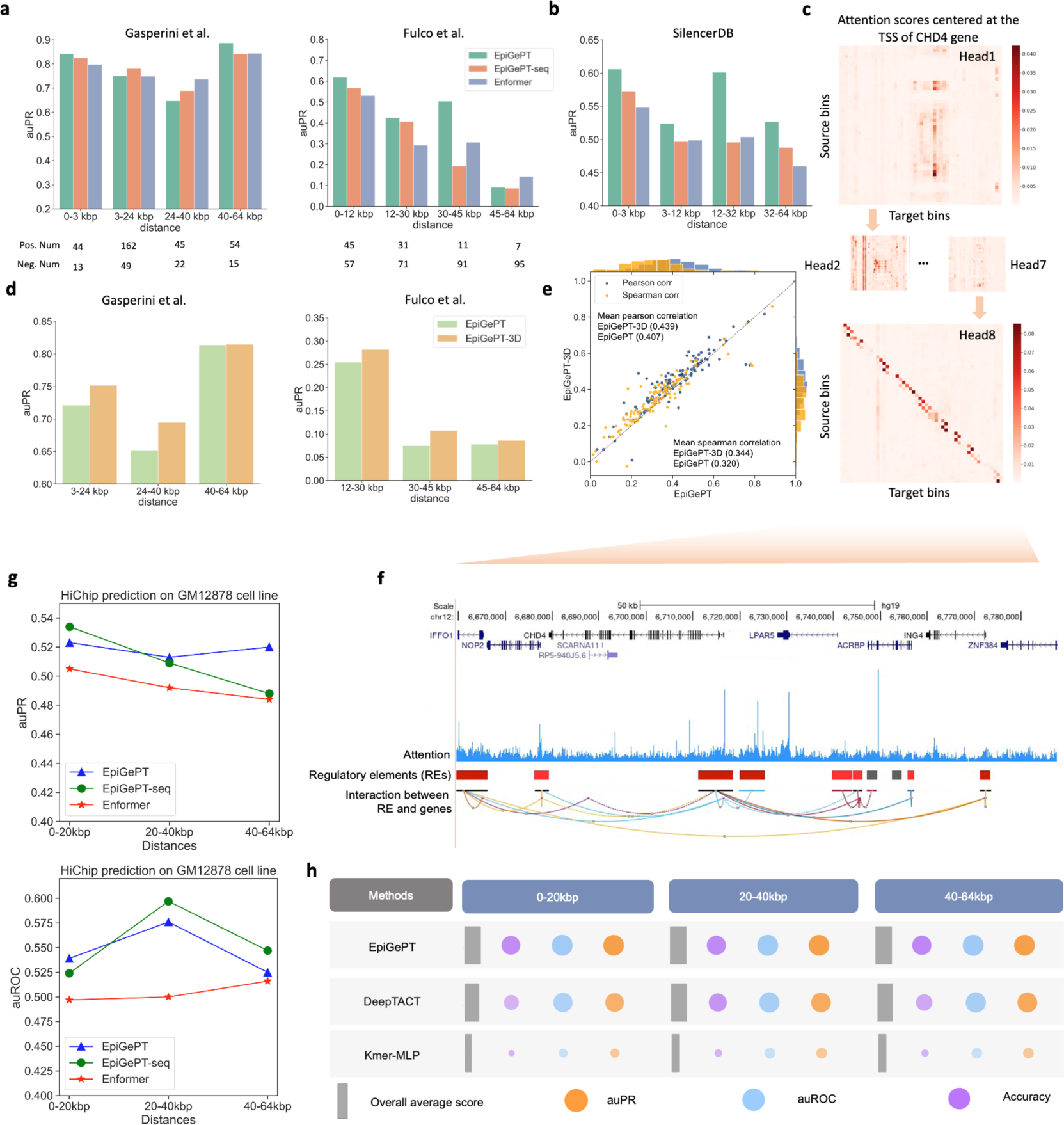
Application of self-attention mechanism in EpiGePT for long-range chromatin interaction identification. **a**, The performance (auPRC) of attention score of EpiGePT in distinguishing enhancer-gene pairs at different distance ranges on two different datasets. **b**, The performance (auPRC) of attention score of EpiGePT in distinguishing silencer-gene pairs at different distance ranges based on the data from SilencerDB^24^. **c**, Heatmap of the self-attention matrix of each attention head centered at the TSS of the *CHD4* gene, the (*i, j*) element in the matrix denotes the average attention score between the *i*th genomic bin and the *j*th genomic bin across all layers. **d**, The performance (auPR) of self-attention scores of EpiGePT and EpiGePT-3D in identifying enhancer-promoter interactions across different distance ranges on the K562 cell type. **e**, The predictive performance (blue points denote pearson correlation coefficients and orange points denote spearman correlation coefficients) of EpiGePT with knowledge guidance across 19 cell types and 15,870 long sequences (128kbp). **f**, Attention scores centered at the TSS of the *CHD4* gene, and putative enhancer regions in its vicinity. **g**, The performance (auROC and auPR) of attention score of EpiGePT in distinguishing HiChIP loops of H3K27ac at different distance ranges on GM12878 cell line. **h**, The performance (auROC and auPRC) of the fine-tuned EpiGePT model and baseline methods (DeepTACT and Kmer) in distinguishing enhancer-gene pairs at various distance ranges (0-20 kbp, 20-40 kbp and 40-64 kbp) on K562 cell line under a 5-fold cross validation setting. The size of the bubbles in the plot represents the magnitude of the metric values, while the width of the gray rectangles along the x-axis signifies the overall average values of the three metrics.

To better understand the self-attention mechanism of EpiGePT, we showed the attention weights (averaged across heads) for the bin containing the TSS of the gene *CHD4*. The attention weights were computed based on the pretrained EpiGePT model with K562 cell line as the context of interest. We also display chromatin interactions detected under K562 as well as regulatory elements annotations from the GeneHancer^27^, It is seen that both the interaction data and the regulatory element annotations are consistent with the attention weights learned by EpiGePT (Fig. 3c and Fig. 3f).

Setting (B): Prediction by EpiGePT-3D, which include Hi-C data in its training. The above results suggest that in a good transformer-based genomic language model, the attention weight given by one bin to another bin (within the input region) should be consistent with the strength of 3D interaction between them. Thus, when experimental data on 3D interaction are available, we can leverage this data to improve the learning of the parameters of our genomic language model, by penalizing parameter values that resulted in poor correlation between the attention weights and the interaction data (see Methods). To obtain such training data, we collected 4,107,687 H3K27ac-based HiChIP loops across 13 cell lines or tissues from HiChIPdb^26^, which denote potential E-P interactions. Setting aside loops from K562 cell line as test data, other HiChIP loops are incorporated into the training. The resulting model is denoted as EpiGePT-3D. We found that adding 3D interaction data in the training can lead to a noticeable improvement for cross-cell-type prediction (3.3% higher in PCC) (Fig. 3e). Moreover, EpiGePT-3D demonstrated improved predictive performance on E-P interactions identified by HiChIP loops in new cellular contexts. For instance, the auPRC increased from 0.652 to 0.695 for Gasperini et al.’s dataset, which is on a context not covered by the Hi-C data in the training, in 24-40kbp group when incorporating 3D genome data.

Setting (C): Prediction by fine-tuning pretrained EpiGePT. Fine-tuning is an strategy that transfers the knowledge of a pretrained model to new tasks, which is particularly prevalent in language models such as GPT^28^ and BERT^29^. Here, we explore the performance of fine-tuning given a pretrained EpiGePT model on downstream tasks, such as predicting 3D genome interaction. Specifically, we fixed the weights of a pretrained EpiGePT model and added an additional finetune network for predicting E-P interactions. We compared EpiGePT with finetuning strategy (EpiGePT-finetune) to two baselines, DeepTACT^30^ and a k-mer frequency based method^29^ with HiChIP H3K27ac loops from K562 and GM12878 cell lines (see Methods). The results illustrate that EpiGePT-finetune exhibited a superior classification performance across diverse distance ranges compared to baselines. For example, EpiGePT-finetune achieved an auROC of 0.949, surpassing 0.866 of DeepTACT ^30^ and 0.771 of Kmer by a large margin in the GM12878 cell line within the 20-40kbp distance range (Fig. 3h, Fig. S9 and Fig. S10). This significant improvement demonstrates the power of fine-tuning a base pretrained genomic language model on a downstream task with limited data.

### EpiGePT unveils the regulatory relationships between TFs and target genes

In this section, we further explored the TF module to see whether EpiGePT is able to learn the regulatory relationships between TFs and target genes (TGs). We defined gradient importance scores (GIS) based on the absolute gradient values of predicted epigenomic signals with respect to the expression of a TF in the input TF profile, to rank the TFs for their potential to regulate a given TG (see Methods). Particularly, we use the TF profile of embryonic stem cell (ESC) to specify the context in the EpiGePT model. We selected the important ESC regulator *POU5F1* as the target gene and calculated the GIS for identifying TF-TF interactions (see Methods). Multiple potential regulators for POU5F1 identified by EpiGePT in ESC context are consistent with literatures, such as *ESRRB*-*POU5F1*^31^ (rank 2^nd^), and *ETV5*-*POU5F1*^32^ (rank 5^th^). Next, we focus on *ESRRB* which plays essential role for balancing pluripotency of ESCs^33^. Treating ESRRB as the target gene, our GIS-based ranking identified several key TFs, such as *POU5F1* and *REST*, that have significantly higher ranks than other TFs (Fig. 4a). By using ChIP-seq data of *POU5F1* for validation, we observed significantly higher GIS in bins overlapping with the ChIP-seq data (Fig. S11, *p*-value < 0.00018 under one-sided Mann-Whitney U test). Next, we visualized the TF ranks obtained from eight epigenomic profiles across 1000 bins surrounding the TSS of *ESRRB*. By averaging ranks across these signals and bins among all the 711 TFs, the important ESC regulator *POU5F1* ranks 3 out of 711 (Fig. 4b). We further collected the top 5% of TFs for each bin and conducted gene ontology (GO) enrichment analysis based on these TF coding genes. Interestingly, the GO terms enriched also included biological processes of embryonic cell differentiation and development. However, using the top 5% of TFs with high expression in ESCs resulted in lower significance for biological processes associated with embryonic cell development (Fig. 4c and Fig. S12), which again demonstrates the effectiveness of the GIS-based ranking. Furthermore, we use TF-TG relationships from either ChIP-seq data or external databases as ground truth to validate the TF-TG relationships inferred by EpiGePT. We defined potential TF-target gene pairs based on TF ChIP-seq data specific to certain cell types among all human genes (see Methods). The results demonstrated a significant higher rank of TF-target gene pairs, compared to TF-non-target gene pairs based on the integrated GIS-based ranking (Fig. 4d, *p*-value < 0.001 under one-sided Mann-Whitney U test). Second, we collected TF-TG regulatory network data from two publicly available databases. We obtained a total of 1,066 TF-TG pairs from the GRNdb^34^ database based on liver-specific GTEx data, and 2,705 TF-TG pairs from the TRRUST^35^ database after filtering. Then we calculated the rank of each TF based on either integrated GIS or the TF expression value by using the liver expression as the TF reference profile. Interestingly, we found that the median ranking percentile of TFs from TRRUST was 3.1%, significantly higher than the percentile of 20.4% based on expression values (Fig. 4e, *p*-value < 1e-5 under one-sided Wilcoxon signed rank test). with a similar result was obtained using another database GRNdb, where EpiGePT is seen to achieve a median ranking percentile of 6.3%, compared to 36.0% by gene expression value. For instance, *TMEM55B*, which plays a significant role in lysosome movement, and is regulated by sterol response element binding factor 2 (*SREBF2*)^36^. Consistently, GIS ranking identified *SREBF2* as the top-ranked TF associated with *TMEM55B*. Overall, the validation results from both ChIP-seq datasets and external databases support the effectiveness of GIS in identifying context-specific TF-TG relationships.

**Fig. 4.**
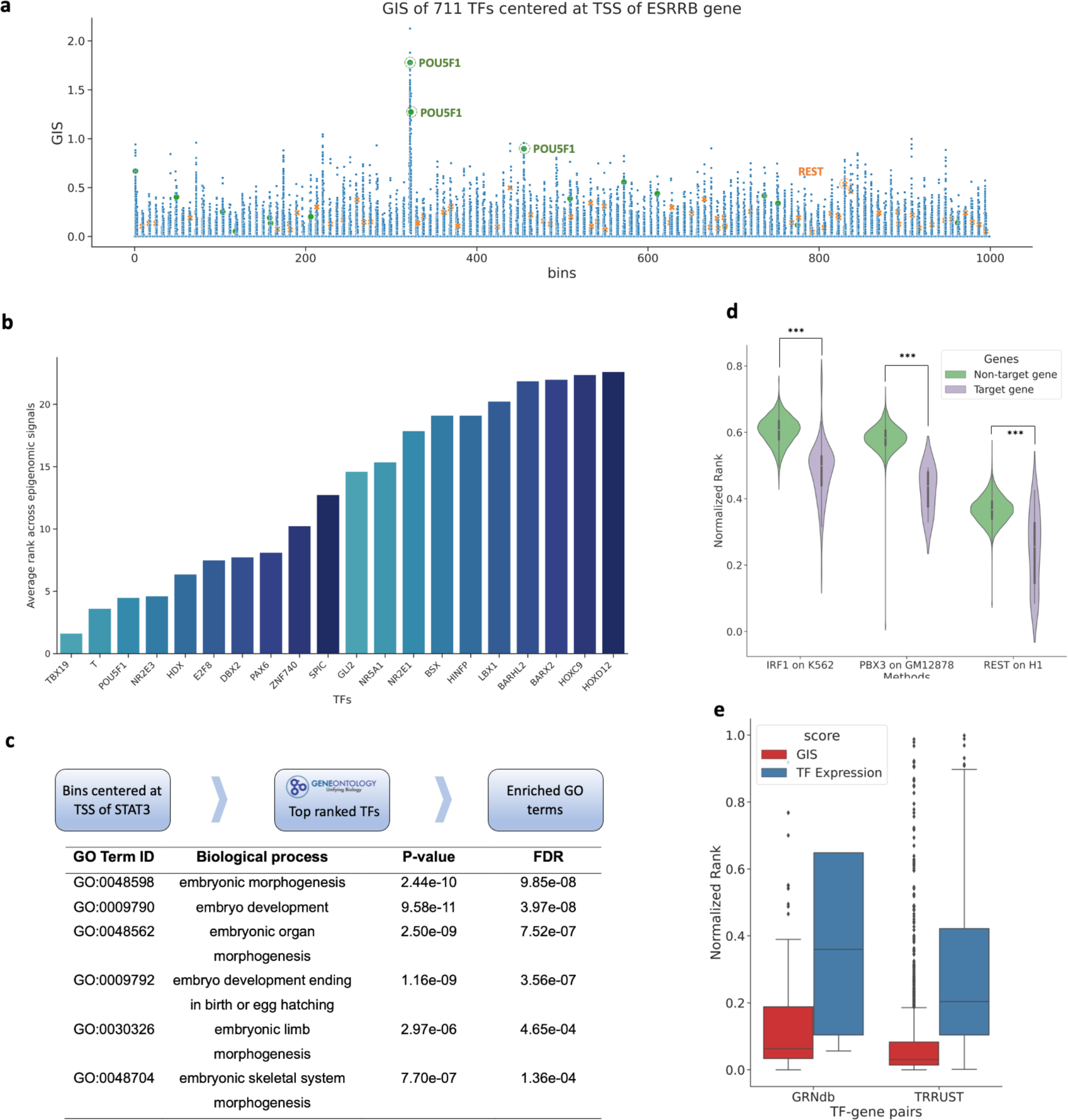
Gradient importance scores (GIS) uncover regulatory transcription factors. **a**, Genomic regions around TSS of the *ESRRB* gene and TF expression data on ESC were used in EpiGePT. The scatter plot represents the GIS scores of 711 TFs on each genomic bin. Each dot represents the GIS score of a core TF on a specific genomic bin. Two important ESC regulators *REST* and *POU1F5* are highlighted. **b**, Bar plot of the top 5% ranked TFs, based on the average ranks from the GIS of eight epigenomic signals across bins (below). **c**, Based on the top 5% ranked TFs in 128kbp region centered at TSS of the *ESRRB* gene, gene ontology enrichment analysis revealed significant enrichment in biological processes related to embryonic development and cellular differentiation. **d**, Based on TF ChIP-seq data, all 23,635 human genes were classified into target genes and non-target genes. The results revealed that TFs exhibited significantly higher ranks on potential target genes compared to non-target genes. **e**, The distribution of the rank of TFs in the GIS and expression value among the 2,705 TF-gene pairs from the TRRUST database and 1,066 TF-gene pairs derived from genotype-tissue expression (GTEx) data of the liver sourced from the GRNdb database.

### EpiGePT improves variant effect prediction

Context-specific prediction of the functional impact of genetic variants is important for genetic studies. To test the utility of EpiGePT in this task, we first collected an eQTLs dataset^37^ that contains 20,913 causal and non-causal variant-gene pairs across 49 different tissues from the supplementary data of Wang et al^37^. EpiGePT, EpiGePT-seq (i.e. EpiGePT without the TF module) and Enformer were then applied to estimate the context-specific log-ratio scores (LOS) between the alternative DNA sequence and the reference DNA sequence, (see Methods, Fig. 5a). Finally, a random forest classifier is trained based on these LOS’s to distinguish causal variant-gene pairs from non-causal pairs. The experimental results show that better prediction performance can be achieved when the LOS is based on EpiGePT than when the LOS is based on Enformer. For example, in the lung tissue, EpiGePT achieved an auPRC of 0.922, compared to 0.873 of Enformer, for the classification of casual SNPs vs non-causal SNPs. To verify the effectiveness of TF module, we replace the TF reference profile of lung with a less relevant cell type, stomach, and the auPRC decreases from 0.922 to 0.892 (Fig. 5b). Similar results were seen for other tissue contexts—across 48 tissues, EpiGePT-seq achieved an average auPRC of 0.910, compared to 0.898 of Enformer (Fig. S4d). The above experiments demonstrated the usefulness of EpiGePT in assessing variant effects.

**Fig. 5.**
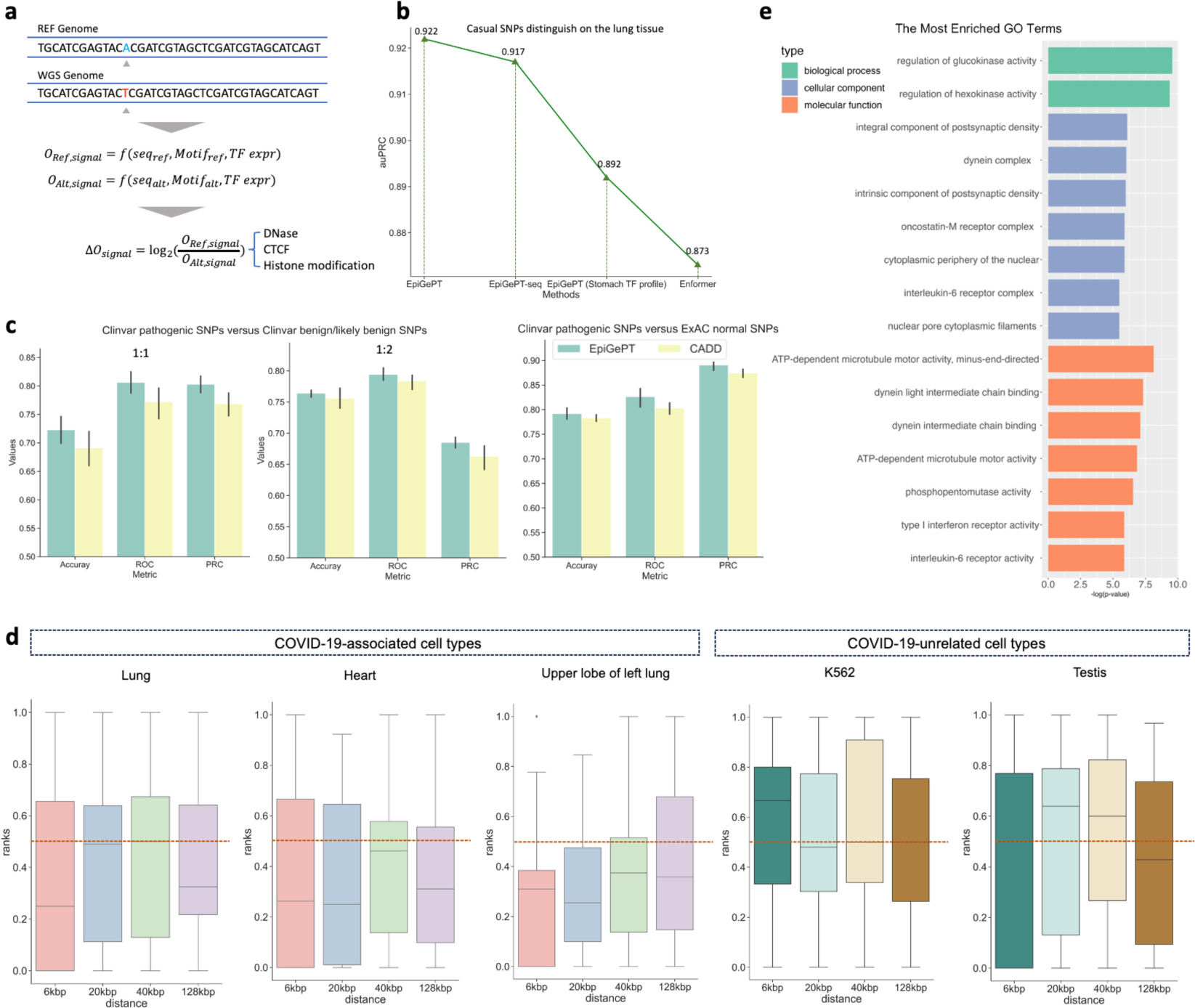
Variant effect prediction of EpiGePT. **a**, The LOS for each epigenomic signal is calculated by the log change fold of the predicted epigenomic signal for reference genome and WGS genome. **b**, The performance of EpiGePT and Enformer in discriminating causal SNPs on the Lung tissue. **c**, The three subplots from left to right respectively depict the classification results for disease-related SNPs and benign SNPs down-sampled sourced from the ClinVar database, with balanced positive and negative samples (1:1 and 1:2 ratio), as well as normal SNPs sourced from the ExAC database with a MLP classifier. **d**, The ranked position of COVID-19 related GWAS data among surrounding benign SNPs based on their LOS, as determined using different tissue or cell-type expression data. The results were stratified based on the distance range of the risk region. The resulting mean and median ranks were both below 0.5. **e**, Enrichment result (Biological process, Cellular component and Molecular function) of the nearest genes of the COVID-19 associated SNPs with the max LOS.

To further evaluate the performance of EpiGePT in predicting disease-associated variants, we extracted 52, 876 pathogenic SNPs from the ClinVar^38^ database and 418, 863 benign SNPs from the ClinVar database, also with 84, 095 benign SNPs from the ExAC database^39^ as positive and negative sets, respectively. We defined a 128kbp region surrounding each pathogenic SNP as the risk region. We extracted all benign or likely benign SNPs that fall within the risk region as the positive samples. As the relevant tissue or cell type information is not available, we concatenated the LOS of the eight epigenomic signals and also the self-attention scores, across multiple cellular contexts, and then evaluated whether the constructed features are beneficial in distinguishing pathogenic SNPs from benign ones in a classification analysis. To achieve this, we augmented the popular CADD-derived features (CADD^40^ scores) by concatenating them to the EpiGePT-derived features discussed in the above, to obtain a comprehensive feature vector (see Methods). Subsequently, we compared the performance of the multi-layer perceptron (MLP) classifier based on the comprehensive feature vector to that based on CADD-derived features alone. The results demonstrated that incorporating EpiGePT-derived features significantly enhance the performance in predicting pathogenic SNPs. Specifically, when the positive-to-negative sample ratio was set to be 1:1, the average auROC increased from 0.772 to 0.806, and the average accuracy increased from 0.690 to 0.723 (Fig. 5c). This observation indicates that features extracted by EpiGePT provide a valuable complement to CADD scores, enabling a more comprehensive interpretation of disease-associated variants.

### EpiGePT prioritizes potential SNPs associated with comorbidities of COVID-19

We investigated whether using EpiGePT to predict variant effects could help in the discovery of key SNPs related to COVID-19. COVID-19 is an infectious disease caused by the SARS-CoV-2 virus, which emerged in late 2019 and quickly spread around the world, causing a global pandemic^41^. In order to validate the ability of EpiGePT in identifying key SNPs, we collected GWAS data from a COVID-19 genetic study^42^, including 9,484 variants derived from 4,933 patients with confirmed severe respiratory symptoms and 1,398,672 control individuals without COVID-19 symptoms. To validate the ability of the model to identify COVID-19-associated SNPs, we firstly defined a risk region around the selected COVID-19-associated SNPs and computed the rank of the variant score of pathogenic SNPs within the surrounding benign SNPs from the ClinVar database. Note that the expected percentile rank for random guessing (uniform distribution) is 0.5 (see Methods). Previous studies^43, 44^ suggested that COVID-19 infection could potentially impair the function of the heart or the lungs, leading to congestive heart failure or decreased lung function. Interestingly, we found that the average rank of COVID-19-associated SNPs was 0.250 when lung expression data was employed for the TF reference profile and a 6-kbp risk region was examined (Fig. 5d, *p*-value < 0.05 under one-sided Binomial exact test). However, when we employed the expression data from less relevant contexts, such as K562 cells or Testis cells, the median rank is close to random guessing (i.e. 0.5), indicating its ineffectiveness in discerning SNPs pertinent to COVID-19. These results suggest that EpiGePT model is able to prioritize the COVID-19-associated SNPs thus shedding lights on finding the potential disease-associated variants and the relevant tissue contexts with our pretrained large language model.

Next, we examine whether the genes close to max-LOS SNPs are likely be associated with biological processes and functions relevant to COVID-19, when compared with genes close to low scores SNPs or not closed to associated SNPs. Since the genetic pathology of COVID-19 is not yet clear and the earliest lesion is in the lungs, we ranked all 9,484 possible SNPs using lung expression data as the TF reference profile. We then identified the SNPs with the highest ranks and performed GO enrichment analysis on nearest genes of the top-30 scored SNPs (Fig. 5e). The enrichment results revealed potential biological processes that are relevant to COVID-19, such as the regulation of glucokinase activity which is associated with the homeostasis of human blood glucose^45^. Notably, diabetes mellitus, a condition closely associated with hyperglycemia, is a typical comorbidity of COVID-19^46^. However, GO enrichment analysis based on the nearest genes of the lowest-scored 30 SNPs resulted in enrichment outcomes that were less relevant to COVID-19 or its complications (Supplementary Fig. S14). Among the potential genes around the top-10 scored SNPs, we identified that the *TBC1D4* gene, which regulates glucose homeostasis, is potentially associated with COVID-19 comorbidities. Our findings are consistent with previous research by Pellegrina et al.^47^ and highlights the potential of our EpiGePT approach in discovering new genetic markers that may be implicated in the pathogenesis of COVID-19. Overall, our EpiGePT model provides new perspectives for understanding how the genetic variants could contribute to the COVID-19 susceptibility and severity.

### Model ablation analysis

To verify the roles of the main modules in the model, we conducted ablation experiments on the model architecture (Fig. S5). For TF module ablation, the results compared to EpiGePT without TF module (EpiGePT-seq) and the inclusion of the TF module led to improvement in cross-cell-type prediction of DNase signals, with a median PCC of 0.787 of EpiGePT, compared to 0.74 for EpiGePT-seq. We additionally examined the impact of the TF module by employing three methods, namely replacing TF scores with zero, replacing TF scores with random noise, and removing motif binding scores. The results again confirmed the positive impact of the TF module (Fig. S5a). For sequence module ablation, we trained a TF-only model without the sequence module. The results indicated that removing the sequence module resulted in an average decrease of 0.084 in the PCCs of the epigenomic signals on a cell-type wise basis (Fig. S5a). For multi-task module ablation, we trained eight separate predictive models for each of the eight epigenomic signals. In the case of the H3K4me1 signal prediction, the performance of the single-task prediction model exhibited an average PCC decrease from 0.408 to 0.329 compared to the multi-task prediction model. Similarly, the overall prediction performance for the eight signals declined by 0.074 (Fig. S5b). This decrease may be attributed to the intricate nature of gene regulation that multiple epigenomic signals can synergize with each other, allowing their joint modeling to gain deeper biological insights.

### Online prediction tool for EpiGePT

In order to facilitate the utilization of EpiGePT for the prediction of multiple chromatin states of any cellular context and genomic regions, we have developed a user-friendly web server, named EpiGePT-online (http://health.tsinghua.edu.cn/epigept/) (Supplementary Text S2). The web server was developed using PHP, JavaScript and HTML, which provides an interactive web interface for efficiently online prediction of epigenomic profiles (Fig. 6). The web server includes a built-in kernel that encompasses the framework for data preprocessing, TF motif binding scores calculating, and prediction of epigenomic signals for both hg19 and hg38 human reference genome. Users can obtain the predicted signals for multiple genomic regions by submitting a region file and a TF expression file in Numpy or CSV formats (Supplementary Table S5), or predicted signals for a specific region by submitting a TF expression file (Fig. S13). We provided TF expression profile across more than 100 cellular contexts from ENCODE on the download page. Users can download the results in csv format for further applications such as genetics analysis. Furthermore, we provide a case application of the EpiGePT-online to enable users to quickly learn how to use our website (Supplementary Text S3). We anticipate that this web server will assist researchers in deepening their understanding of gene regulatory mechanisms.

**Fig. 6.**
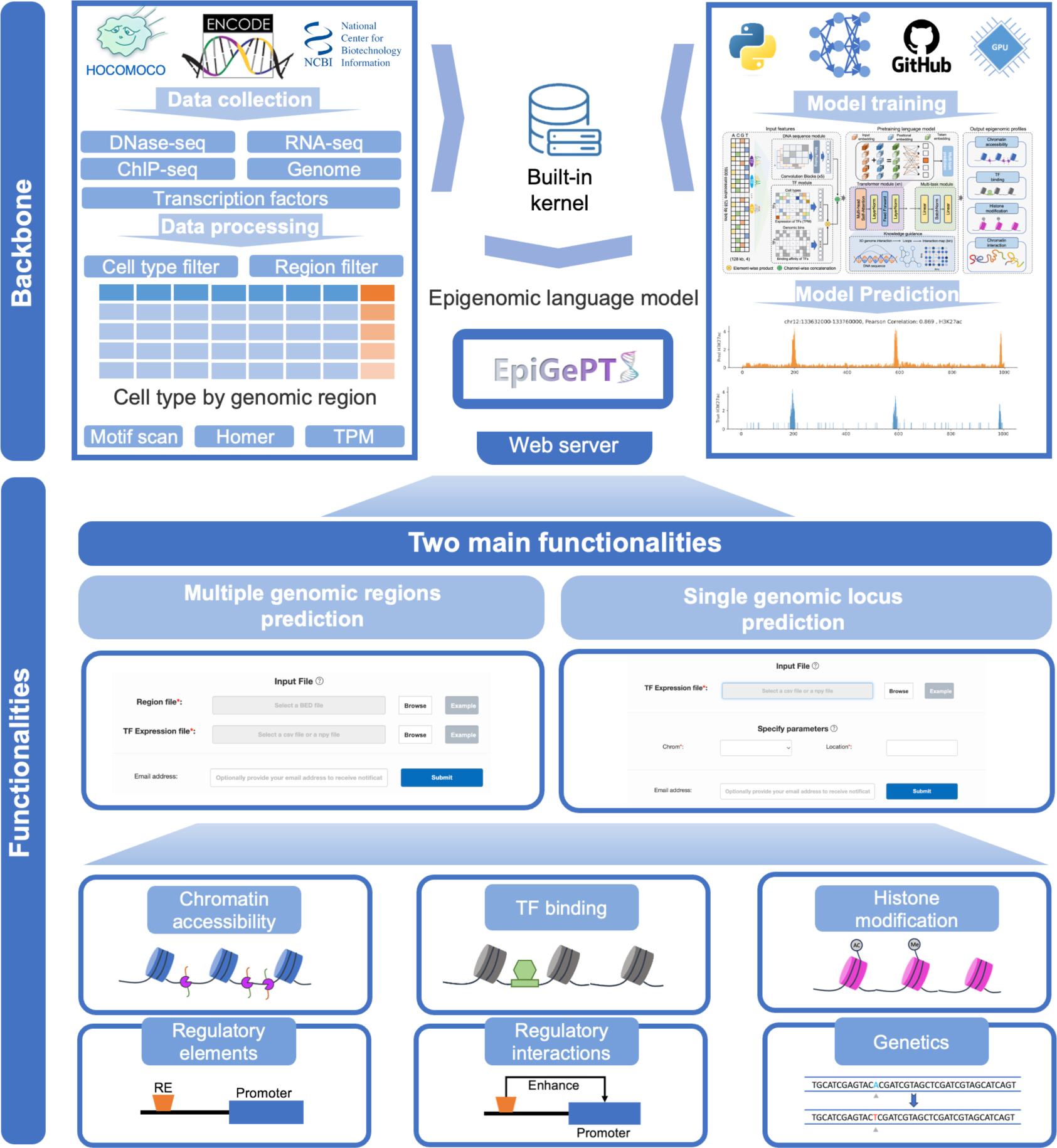
Overview of the online prediction web server of EpiGePT. We collected eight types of epigenetic genome modification signals and corresponding expression data of transcription factors in different cell types or tissues from the ENCODE project. Based on these data, we trained the EpiGePT model and deployed it as a built-in kernel on an Apache server. Users without much coding experience can also access the web server in two ways to obtain the eight types of epigenetic genome modification signals for specified cell types and genomic regions without programming or installation.

## Discussion

In this paper, we introduced a pretrained transformer-based language model for epigenomics. Compared with the existing task-specific models and sequence-based language model, EpiGePT has the added capability to make predictions on novel contexts. Furthermore, EpiGePT is able to incorporate a new type of data (3D genome interaction data) during model training, which enables the identifying functional regulatory interactions such as enhancer-promoter interactions. EpiGePT demonstrates state-of-art performance in diverse experimental settings compared to existing methods. Based on the predicted epigenomic features and 3D interactions from EpiGePT, we performed two investigations on how information is encoded in the human genome sequence: First we identify the interactions of cis-regulatory elements and their target genes with the help of self-attention mechanism in EpiGePT. Through direct utilization of self-attention scores, model fine-tuning, and leveraging 3D genome interactions, we validated the capacity of EpiGePT to capture regulatory interactions. Second, to assist the identification and interpretation of human disease-associated SNPs, we estimate the effect of a variant on the epigenomic features around the variant, based on the LOS computed by the outputs of EpiGePT. Such variant effect prediction by EpiGePT establishes a foundation for understanding the underlying relationship between genetic variations and disease mechanisms.

There exist several extensions and refinements that can be applied to further improve the EpiGePT model. Firstly, the incorporation of chromatin regulators (CRs) as trans-acting factors into the TF module could enhance the modeling of regulated transcription processes, thereby increasing the accuracy of the predictions. Second, the integration of DNA methylation information^48^ while modeling DNA sequences allows for a more comprehensive and accurate decoding of the epigenomic language, providing a more comprehensive model of gene regulation states compared to the analysis solely based on DNA sequences. Third, the rapid advancements in sequencing technologies have enabled the accumulation of vast amounts of multi-omics data, encompassing different scales from biomolecules to single cells, tissues, and organs^49^. The integration of multiscale and multi-omics information is a trend and a major challenge in deciphering gene regulatory landscapes. Integrating single-cell level data into EpiGePT is an important direction for future improvement. For example, utilizing clustered single-cell multi-omics data as pseudo-bulk data can further expand the training context of EpiGePT. The application of EpiGePT to single-cell epigenomics could enable the profiling of chromatin signals at single-cell resolution, facilitating a holistic understanding of regulatory heterogeneity in different cell subpopulations.

Based on EpiGePT, users are able to predict multiple chromatin profiles in different cell lines or tissues, which could provide a foundation for biological discovery, decoding transcriptional regulation mechanisms, and investigating disease mechanisms. We anticipate EpiGePT will furnish researchers with valuable insights into understanding regulatory mechanisms.

## Methods

### Data processing

#### Chromatin accessibility data and Expression data

We used three different datasets in the experiments. For chromatin accessible data, we downloaded DNase bam files and narrow peaks across 129 human biosamples from ENCODE^19^ project (Supplementary table S1 and S2). We divided the human hg19 genome into 200bp non-overlapping bins, and we assigned the label for each bin in each cell type. For the regression design, we pooled the bam files of multiple replicates for a cell type (Supplementary table S1 and S2), and obtain the raw read count *n*_*lk*_for bin *l* in cell type *k*. We normalized the raw read count in order to eliminate the effect of sequencing depths, in the form of *ñ*_*lk*_ = *Nn*_*lk*_/*N*_*k*_, where *N*_*k*_ denotes the total number of pooled reads for cell type k and 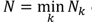 denotes the minimal number of pooled reads across all cell types. The normalized read counts are further log transformed with pseudo count 1, which represent the continuous level of chromatin accessibility. For binary classification design, we assigned a binary label *y*_*lk*_ to 1 if the number of raw read counts of the bin *l* in the cell type *k* greater than 30, which represent the bin is an accessible region in this cell type, resulting in the identification of regions as accessible in 13% on average and 8% at median in the screened genomic regions across 129 cell types. The proportion of open regions varies among different cell types, and the average openness level mentioned above is generally consistent with that maintained in ChromDragoNN^6^.

RNA-seq data of the 711 human transcription factors were downloaded and extracted from the ENCODE project (Supplementary table S5 and S6). We perform log transformation with pseudo count 1 and quantile normalization based on TPM values. The normalized TPM values were averaged across replicates and mean expression profile after normalization of each cell type was finally used to calculated of the transcription feature.

#### Multiple chromatin signals data

For the human reference genome hg19 (GRCh37), DNase-seq, RNA-seq and ChIP-seq data were also downloaded from ENCODE project (Supplementary table S3, S4 and S6). We applied the same process to these data as above, and finally we obtained the 8 epigenomic signals of 13,300,000 bins of 128bp in 28 cell types. The continuous level of chromatin signals we extracted were ‘DNase’, ‘CTCF’, ‘H3K27ac’, ‘H3K4me3’, ‘H3K36me3’, ‘H3K27me3’, ‘H3K9me3’ and ‘H3K4me1’, which includes crucial epigenetic modifications and markers for gene regulation and transcription.

For the collected the data of human reference genome hg38 (GRCh38), we adopted a data collection strategy that includes missing data. Specifically, within a particular tissue or cell type, we ensured the presence of at least one ChIP-seq signal. Then, epigenomic profiles of 8 signals for 15,870,000 bins of 128bp across 104 cell types were obtained.

### Model architecture

#### Sequence module

As shown in Fig. 1 and Fig. S2a, the sequence module receives a one-hot matrix (A = [0,0,0,1], C = [0,1,0,0], G = [0,0,1,0], T = [0,0,0,1]) of size (128000,4) as input, representing a sequence of 128 kilobase pairs (kbps) and contains five 1-dimentional convolutional blocks to extract DNA sequence features. Each block includes a convolutional layer and a maxpooling layer (Fig. S2b). The first convolutional layer considers the input channels as 4 and performs convolution along the sequence direction. The input sequence features are one-hot embeddings of size *L* × 4, where *L* denotes the length of the input long range DNA sequence. After 5 maxpooling layers, the output size of sequence feature is *L*/*N* × *C*, where *C* denotes the hyper-parameter for sequence embedding and N denotes the length of locus to predict. We set C to 256 in the pre-training stage of chromatin accessibility prediction experiments. Rectified linear units (ReLU) are used after each convolution operation for keeping positive activations and setting negative activation values to zeros. By reducing the input length by 128 times through pooling operations, this module effectively compresses the input information while retaining essential features. Sequence features were then concatenated with TF expression features, and we finally obtained a vector of size *L*/*N* × (*C* + *n*_*TF*_), where *n*_*TF*_denotes the dimension of the transcription factors features after padding. In our model, after adding padding to the 711 TFs, the *n*_*TF*_is set to 712. Therefore, the input token number for the transformer module is 1000, and each token embedding has a dimensionality of 968.

#### Transformer module

We utilize the transformer module to integrate information from both the sequence and transcription factors (TFs), enabling the capturing of long-range interactions between genomic bins. We applied *N*_*t*_layers of Transformer encoder with *n*_ℎ_different attention heads to the token embedding sequence. The input word embedding (*X)* of the transformer encoder is a genomic bin sequence with dimensions (*Sequence length, embedding dim*). Specifically, this dimension is (1000, 968) in EpiGePT, indicating that input genomic bin sequence has a length of 1000, and each genomic bin has an embedded representation that combines the sequence information with cell-type-specific features with dimension of 968. For position embedding, we employed absolute position embedding to represent the positional information of the 1000 genomic bins in the input 128kbp DNA sequence, with dimensions of (1000, 968). Each Transformer encoder includes a multi-head self-attention mechanism and a feed-forward neural network. For self-attention in each head, the calculation is based on the matrix operation.

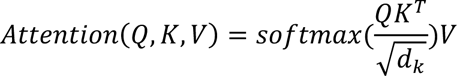

For multi-head attention, Transformer encoder learns parameter matrices 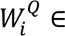 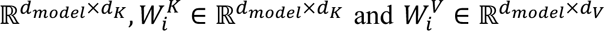 for the *i*_*t*ℎ_ head and concatenate the multiple heads to do the projection, then learns parameter matrices *W*^0^ ∈ ℝ^*n*_h_d_v_×d_model_^ to obtain the output of multi-head attention layer.

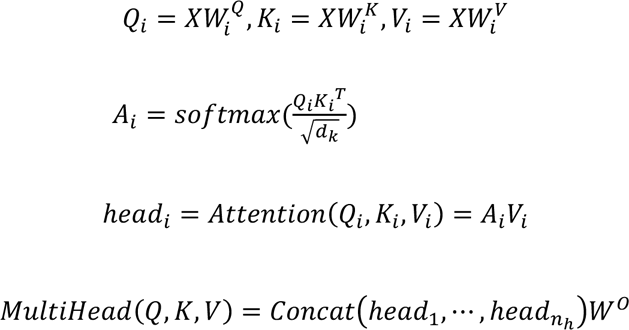

where *d*_*model*_denotes the dimension of token embedding *X*, which is 968 in EpiGePT *X* denotes the embeddings from the sequence module for the first attention layer or the output of previous attention layer. *n*_ℎ_denotes the number of head in Transformer encoder, which is 8 in EpiGePT, and *d*_*K*_ = *d*_*V*_ = *d*_*model*_/*n*_ℎ_ = 121. The matrix *A*_*i*_ is called the self-attention matrix for head *i*. The outputs of *n*_ℎ_ heads are then concatenated, and a mapping function represented by *W*^-^ is applied to obtain the output of the multi-head attention. After passing through an add & norm layer, the multi-head attention output is used as input to the feed-forward layer, where more comprehensive features of the input sequence are extracted. The above describes the computational workflow of a single Transformer encoder layer. We set *N*_*t*_to 16 for the chromatin accessible prediction experiments, *N*_*t*_ to 12 for the chromatin state classification and multiple chromatin signals prediction experiments.

#### Prediction Module

For regression model, the output layer uses a linear transformation and use mean square error (MSE) as the loss function. For classification model, the output layer uses a linear transformation combined with a sigmoid function, and use the cross-entropy loss for classification experiments.

#### TF module

For binding status, we scanned the input bins for potential binding sites for a set of 711 human transcription factors from HOCOMOCO database^50^ with the tool Homer^51^ (Table S5). We then selected the maximum score of reported binding status for each transcription factor to obtain a vector of 711 dimensions as the motif feature for each DNA bin. For gene expression, we focused on log-transformed TPM values of the 711 transcription factors and obtained a vector of 711 dimensions after quantile normalization as the expression feature. With these data, we combined the two vectors of motif and expression features by taking the element-wise product, and we concatenated the result to the output of sequence module.

### Model evaluation

To evaluate our model, we applied five-fold cross-validation in the different experiments on cell-type level. For chromatin accessible experiments, the 129 cell lines are partitioned into a training set and a testing set randomly.

Cell-type-wise metrics are defined to evaluate our method in different experiments, which were calculate with the data within a test cell type across all genomic locus. For binary classification design, we used cell-type-wise auPRC and auROC to evaluate our EpiGePT. Let *Y*_*L*×*k*_ and *Ŷ*_*L*×*k*_ be the true and predicted matrix, where L denotes the number of locus and K denotes the number of test cell types. We calculated the auPRC and auROC for each (*y*_1*i*_, *y*_2*i*_, ⋯, *y*_*Li*_) and (*ŷ*_1*i*_, *ŷ*_2*i*_, ⋯, *ŷ*_*Li*_). For multiple classification, we use macro average of the auPRC and auROC to evaluate the classification performance, which compute the metric independently for each class and then take the average hence treating all classes equally. For regression design, we used two metrics for model evaluation, which are cell-type-wise Pearson correlation coefficient and prediction squared error. Prediction square error (PSR) is calculated as *PSR* = 1 −∑_*k*_ ∑_*l*_(*y*_*lk*_ − *ŷ*_*lk*_)^2^/(*y*_*lk*_ − *ȳ*_∗*k*_)^2^, where *ȳ*_∗*k*_ = ∑_*l*_ *y*_*lk*_/*L* denotes the mean of the true level of the response in the cell type k.

To compare the performance of our method with other baseline methods, we conducted hypothesis testing on the metrics based on cell types. Since the metrics on a given cell type across different methods are paired data and the statistical distribution is unknown, we employed both Binomial and Wilcoxon tests, with the alternative hypothesis being that EpiGePT outperforms the other methods. If we reject the null hypothesis, it provides compelling evidence to support the claim that EpiGePT performs better than the other methods.

To evaluate the computational efficiency, we recorded the running time of a single epoch of EpiGePT and baseline methods (Supplementary Text S4). Compared to traditional CNN models such as DeepCAGE^16^ and ChromDragoNN^6^, as well as larger sequence models like Enformer, EpiGePT demonstrates a balance between high computational efficiency and performance.

### Model training strategy

As our proposed model is designed for cross-cell-type prediction of epigenomic signals by multi-task learning. Some of the target epigenomic signals are missing in the existing ENCODE database. For instance, there are 104 cellular contexts with both gene expression and at least one of the epigenomic data. However, this number will decrease from 104 to 28 if we consider eight epigenomic signals simultaneously. The proposed model takes each cellular context and genomic region pair as a training instance, which ensures the availability of a very large number of training instances. To utilize the data from the cellular contexts where some signals are not available (missing data), we will use a new training strategy to handle the missing data where the loss function is designed as

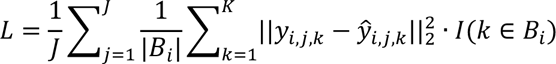

where *y*_*i*,*j*,*k*_ and *ŷ*_*i*,*j*,*k*_ denote the *k*^*t*ℎ^ true and predicted signal from the *j*^*t*ℎ^ genomic bin in the *i*^*t*ℎ^ context, and *I*(·) is an indicator function and *B*_*i*_ denotes the index set that contains all available signals in the *i*^*t*ℎ^context. We update the parameters in the model through stochastic gradient descent based on minibatches. We utilized the Adam optimizer with a batch size of 10 and a learning rate set to 5 × 10^−5^. This training strategy provide us with a significantly larger training sample size and allows us to utilize much more available data from the public databases, and we enable EpiGePT to learn broader patterns of epigenetic states across diverse cell types.

### Incorporation of 3D chromatin interaction data

With the emergence of methodologies like Hi-C and HiChIP for genome-wide chromatin interaction measurement, a substantial volume of 3D chromatin interaction data has been produced across various cellular contexts. Clearly, this data can provide highly valuable information for identifying functional elements in the genome and for understanding gene regulation, but this information has not been captured by current genomic LLMs such as the Enformer^15^ or earlier CNN-based genomic models^6, 7, 16, 52^.

We propose here to exploit the self-attention weights of the transformer model to design a learning strategy that would allow EpiGePT to capture interaction information from Hi-C or HiChIP data. Specifically, we propose to use the ground truth 3D genome interaction to guide the self-attention matrices in the transformer module during the training process. First, we obtained loop information at 5k resolution from the HiChIPdb database^26^. Given potential noise within HiChip data, we selectively filtered potential H3K27ac-based HiChip loops using a stringent q-value threshold of 0.001. This curation aimed to utilization of highly confident loops, safeguarding the model’s ability to capture regulatory information without interference from noise. In this way, we acquired corresponding HiChip loop data for 13 out of 104 cell types. Next, we mapped these loops onto the genomic bins used for pre-training. Specifically, we employed the normalized count as a metric to gauge the likelihood score for each loop. During the mapping process, we aggregated all loops based on this score to each specific genomic bin, and then we obtained the HiChIP interaction matrix *H*_*i*_. Based on the self-attention matrix 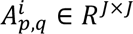 and the HiChIP interaction matrix *H* from the *i*^*t*ℎ^ cell type/tissue where *p*, *q* are indexes for transformer layer and multi-heads, we apply a row-wise normalization to *H*_*i*_(row sum to 1) to obtain 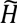_*i*_ and average the self-attention matrices across the heads in the last transformer layers to obtain 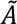^*i*^.. Since elevated attention weights are expected between regions that interacts in 3D, we will compute a new loss term CSL, which is defined as cosine similarity loss between the rows of 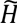_*i*_ and 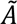^*i*^. Through the guidance of 3D genome interaction data, our approach can learn a more comprehensive model for gene regulation. For example, it will enable prediction of cell-type specific enhancer-promoter interaction, which is a task beyond current models such as the Enformer. Note that the CSL term does not alter the architecture of the model. It simply put some soft constraints on the attention weights according to the experimental data on chromatin interactions, so that the optimized model will give predictions that are more consistent with the context-specific interaction data. During training, the weight α for 3d genome loss was chosen as 2.

### Fine-tunning for predicting E-P interaction

For the fine-tuning process, we kept the parameters of the pre-trained model fixed without making any updates. For the specific fine-tuning task of chromatin interaction prediction based on HiChIP data, the multi-task prediction module was replaced with a two-layer MLP network, containing 256 hidden nodes for each layer. During the training process, only the weights in the MLP network in the prediction module were updated. Notably, when utilizing HiChIP data at a resolution of 5k, both the enhancer and promoter anchors spanned 5kbp. Then we use a region extending 128kbp from the center of the anchor of the neighboring gene, as input region for EpiGePT. Consequently, a 968-dimensional feature vector for each genomic bin was derived from the output of the last transformer encoder layer. These feature vectors from all bins within the two anchors were concatenated, resulting in a high-dimensional vector of size 76,472. To ensure the fairness of validating EpiGePT-finetune in capturing E-P interaction relationships, we fine-tuned the model separately on the HiChIP data of each cell line during the fine-tuning process. The test cell lines K562 and GM12878were excluded from the pretrained EpiGePT training cell types.

### Baseline methods

Four baselines were introduced for epigenetic signals prediction. BIRD^17^ is a multiple linear regression model that only takes gene expression data as input and makes predictions on a fixed locus. ChromDragoNN^6^ is a deep neural network that takes gene expression of 1630 TFs and DNA sequence as input. Specifically, ChromDragoNN^6^ uses a ResNet^53^ to extract sequence features and use linear transformation to combine the TF gene expression feature and sequence feature to make the final prediction. DeepCAGE^16^ is a deep densely connected convolutional network for predicting chromatin accessibility. Enformer^15^ is a deep neural network that integrates convolutional neural network and transformer, and only takes DNA sequence as input. Enformer takes DNA sequence of length 196kbp as input to predict 5,313 genomic tracks of human and 1,643 tracks of mouse genome simultaneously. Enformer can only model and predict cell types in the training data and cannot be applied to new cell types. In order to ensure fairness in some of the benchmark experiment, we retrained the Enformer model with the same input and output data as EpiGePT with Pytorch-lightning and made modifications on the number of encoder layers when reproduce the Enformer model (Supplementary Text S5). Besides, comparison with the pretrained Enformer model was also provided in Fig.2d where we strictly used the ENCODE experiment ID to obtain the matched experiments for comparison.

Two baseline methods were introduced for predicting HiChIP interaction. DeepTACT^30^ is a deep learning method for predicting 3D chromatin contacts using both DNA sequence and chromatin accessibility. We adopted the structure of DeepTACT^30^ and kept the anchor length at 5k. The input to the model consists of two anchor sequences represented as one-hot matrices and the two openness scores of the two anchors on the corresponding cell type extracted from OpenAnnotate^54^. Regarding the Kmer features^55^, K is chosen as 5 to extract sequence features. For each anchor, a vector of dimension 4^5^ = 1024 was obtained. Further training was performed using an MLP with a hidden layer dimension of 256.

### Prediction of 3D genome interaction

We collected cis-regulatory elements-gene pairs in K562 cells from other studies and public database to demonstrate the interpretability of self-attention mechanisms in the EpiGePT. Enhancers and silencers are typical *cis*-regulatory elements known play important roles in transcriptional control during normal development and disease. For enhancers, we downloaded enhancer-gene pairs from two studies: Gasperini et al.^22^ and Fulco et al.^23^, both of which were tested using a CRISPRi^21^ assay perturbation. Two datasets contain 664 and 5,091 element-gene interactions. For silencers, we obtained and random sampled 831 validated silencers-gene pairs with distance within 64kbp in K562 cells curated from high-throughput experiments from SilencerDB^24^. As there are no experimentally validated interaction relationships between these silencers and genes, we generated silencer-gene pairs by associating the nearest neighbor genes for classification purposes. Similarly, negative samples were generated by constructing DNase-seq, ATAC-seq and nearest genes using the same approach. Ultimately, we obtained a dataset comprising 1,662 silencer-gene pairs, encompassing both positive and negative instances.

To obtain scores for regulatory element-gene pairs, we first used the region extending 128kbp from the center of the enhancer as input and extracted the token where the interacting genes reside, so that we could filter out regulatory element-gene pairs that were located further than 64kbp apart. Subsequently, we stratified the remaining pairs based on their distance. Since the positive and negative sample ratios varied across datasets, we adopted different stratification strategies for different distance ranges (Fig. 3). Next, we averaged the attention matrices of the Transformer encoder across all layers and heads. The summed attention scores from other tokens to the key token containing the gene TSS were used as the attention score of this element-gene pair. This score represents the attention value that the enhancer-centered region receives for the TSS of the gene. We also calculated the attention score from the bin containing the center of the regulatory element to the bin containing the TSS, which only slightly affects the experimental results of regulatory element prioritization.

We collected 5k resolution data from the HiChIPdb (http://health.tsinghua.edu.cn/hichipdb/) database, specifically from K562 and GM12878 cell lines. We filtered the data to include only loops where at least one anchor falls within a gene region. We stratified the loops based on distance into three categories: 0-20kbp, 20-40kbp, and 40-64kbp. For each distance category, we selected 2000 positive pairs with most significant q-value. To ensure consistency in the distance distribution, we selected negative pairs by fixing a gene and choosing anchors at equidistant locations in the opposite direction. These are then used to as test data to evaluate the prediction methods.

### Gradient importance scores

EpiGePT possesses the capability to assign priority rankings to transcription factors by utilizing gradient importance scores (GIS), taking into account specific cell types and chromatin regions. The GIS were employed to identify potential functional relationships between specific TFs and target genes. Specifically, for a given TF-target gene pair, the TSS of genes were used as central loci, and the regions spanning 128 kbp upstream and downstream of the TSS were selected as input. Next, we selected bins with motif binding scores indicating potential binding for the given TF. For these selected bins, we calculated the GIS for the predictions of eight epigenomic signals, for each of 711 core TFs.

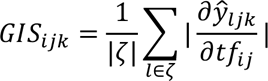

Where, *i* denotes the *i*th TF in the set of core TFs, *j* denotes the *j*th cell type, *k* denotes the *k*th predicted epigenomic signal, and *ζ* denotes the set of genomic bins that have binding for the given TF. In the calculation of the gradient, *ŷ*_*ljk*_denotes the predicted value of the *k*th epigenomic signal by the model using the expression in the *j*th cell type at the *l*th bin. On the other hand, *tf*_*ij*_denotes the product of the expression of *i*th TF in the *j*th cell type and the corresponding TF binding score.

If we consider the GIS for the prediction of all 8 epigenomic signals simultaneously, we can prioritize the TFs by calculating their ranks based on each signal separately. Then, we can calculate an integrated gradient importance score (IGIS) for each TF by averaging the ranks from all 8 signals.

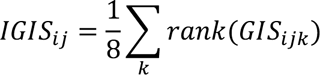

Both the GIS and the IGIS are capable of capturing the significance of a transcription factor (TF) in regulating a specific gene within the context of a specific cell type. Consequently, these scores hold potential value in the discovery of TFs that play crucial roles in the regulation of specific genes, thereby contributing to our understanding of essential regulatory mechanisms.

In the context of validating TF-TG pairs in the GRNdb and TRRUST databases, we opted to utilize liver expression data as a representative example due to the unavailability of cell type information for TRRUST. Furthermore, in this experimental setup, the *tf*_*ij*_denotes the expression of *i*th TF in the *j*th cell type and *ζ* denotes the set of genomic bins that have binding for the TF of the given TF-target gene pair.

### Potential TF-target gene pairs from ChIP-seq data

In this study, we utilized three distinct cell types to conduct a comprehensive screening of TF-target gene pairs and non-target gene pairs across the human genome. Initially, we obtained the narrow peak files (ENCFF388AJH, ENCFF717IXP, and ENCFF885KLR) from ChIP-seq experiments across three cell types from the ENCODE project. Subsequently, we examined the number of peaks within a 128kbp region both upstream and downstream of the TSS for each gene. Different thresholds were applied to the ChIP-seq data of various TFs. Genes lacking any peaks within the defined region were classified as non-target genes, while genes surpassing the threshold in terms of peak counts were designated as target genes. Specifically, for the aforementioned three cell types, threshold values of 10, 15, and 6 were respectively employed. Finally, the IGIS approach was employed to determine the corresponding ranks of TFs in the TF-target gene pairs.

### Pathogenic SNPs prioritization

We collected single nucleotide polymorphisms (SNPs) data from the ClinVar and ExAC databases, which include both potentially pathogenic and benign SNPs. To evaluate the ability of EpiGePT to predict variant effects, we computed the log-ratio scores (LOS) for multiple chromatin signals using EpiGePT on these SNPs. Subsequently, we utilized these scores to distinguish between pathogenic and benign SNPs. The LOS for each chromatin signal was defined by computing a forward pass through the model using the reference and alternative alleles.

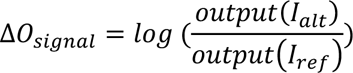

Where *I*_*ref*_denotes the input DNA sequence based on the reference genome, and *I*_*alt*_denotes the input DNA sequence containing variants. Each chromatin epigenomic profile in each cell line or tissue predicted by EpiGePT can be used to compute a specific variant score. We did not take the absolute value in this calculation, so the resulting LOS indicates the direction of change in the model output after the appearance of the variant. In addition to the predicted chromatin signals output by the eight models, attention score changes based on self-attention are also noteworthy. We computed the log-ratio scores for attention by summing the attention scores of the 10 bins upstream and downstream of the locus of the SNP, to evaluate the effect of the variant.

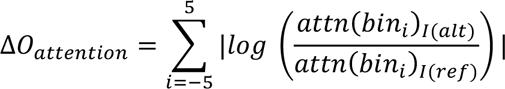

Where *i* represents the index of the neighboring bins relative to the locus of the SNP. To avoid the variant effects of different bins from cancelling each other out during the summation process, we computed the absolute value of the change in attention scores for each bin and then summed the scores of the 10 adjacent bins centered at the SNP position. For the classification of pathogenic SNPs, we calculated these nine LOS for attention separately for each of the 28 tissues or cell lines in training data. As a result, we obtained a feature vector of 252 dimensions for each SNP. Then a classifier with 252 features computed by EpiGePT and 52 annotations from CADD score as inputs are used to predict pathogenic SNPs against benign or likely benign SNPs. Here, we employed MLP as classifier to validate the effectiveness of the features we obtained. A five-fold cross-validation experiment is employed for validation, and we utilize two different positive-to-negative sample ratios, namely 1:1 and 1:2. For each sample ratio, we randomly sample 32,000 positive samples. The effectiveness of the variant score in identifying pathogenic SNPs is evaluated using the area under the auROC and the auPRC. Additionally, we also utilized the logistic regression (LR) as the classifier, consistent with the LR classifier used in CADD, and found a similar improvement when predicting pathogenic SNPs.

### COVID-19-associated SNPS prioritization

We applied the same method to calculate the LOS of the 8 epigenomic signals for the COVID-19 GWAS data. The absolute values of the scores were summed as the overall score for each SNP. Then, we use the absolute sum as the effect score of the SNP and prioritize the COVID-19-associated SNPs based on this score. For each significant SNP associated with COVID-19 severity obtained from the GWAS data, we selected normal SNPs within a 128kb region around the SNP as background to calculate the rank of the LOS for the COVID-19 associated SNP in this region. Furthermore, we calculated the LOS for all 9,484 COVID-19 associated SNPs and ranked them accordingly. The top 10 SNPs with the highest LOS were selected, which are considered to have potential genetic associations with COVID-19 severity and complications.

### GTEx classification

We collected eQTL data from the supplementary materials of Wang et al^37^. In their study, the authors identified causal eQTLs through statistical fine-mapping, using a posterior inclusion probability (PIP) threshold of >0.9 for putative causal variants based on expression modifier score (EMS), and a PIP threshold of <0.9 for putative non-causal variants. To validate the ability of EpiGePT to distinguish potential causal variants, we perform a classification task on these variants. For each variation, 128kbp sequence regions near it were selected as the input of the model, and a score of variation was given by EpiGePT model. For each variant under each tissue, we can obtain an 8-dimensional vector of genomic features including DNase, CTCF and other ChIP-seq signals. Based on the LOS, separate random forest classifiers consisting of 10 decision trees are trained for each tissue in order to distinguish between causal and non-causal variants. The models are evaluated using 5-fold validation on each tissue, with area under the auPRC and auROC as metrics for assessing their ability to distinguish between causal and non-causal variants.

## Code availability

All components of EpiGePT are freely available at https://github.com/ZjGaothu/EpiGePT. Here, users can access the code for reproducing EpiGePT, as well as the data collection and preprocessing pipelines used for model training in benchmark experiments.

## Data availability

Information and processed data of multiple chromatin signals of whole genome, motif binding status and expression data of TFs in the corresponding cell lines/tissues, which are used in EpiGePT are available at Supplementary Tables. The information about the cell lines/tissues used and the 711 filtered transcription factors is available in the supplementary table. The High throughput validated silencers of K562 cell line are download from SilencerDB (http://health.tsinghua.edu.cn/silencerdb) database. The HiChIP data of K562 cell line and GM12878 cell line are downloaded from HiChIPdb (http://health.tsinghua.edu.cn/hichipdb/) database. The DNase-seq peak and ATAC-seq peak data are obtained from the ENCODE project. Enhancer-gene pairs of CRISPRi^23^ experiments are obtained from the supplementary information of Gasperini et al. and Fulco et al. The regulatory network data for transcription factors and target genes were obtained from the TRRUST^35^ database (https://www.grnpedia.org/trrust/) and the GRNdb^34^ database (http://www.grndb.com). The annotated chromatin states for whole genome are downloaded from the ROADMAP epigenomics project (https://egg2.wustl.edu/roadmap/web_portal/chr_state_learning.html). The RNA-seq read counts matrix for protein coding genes used for the prediction of the chromatin 15-states annotated by ChromHMM are downloaded from the ROADMAP project (https://egg2.wustl.edu/roadmap/data/byDataType/rna/expression/57epigenomes.N.pc.gz). The GWAS data of COVID-19 are download from the COVID-19 Host Genetics Initiative (https://www.covid19hg.org/).

## Ethics declarations Competing interests

The authors have declared no competing interests.

## Acknowledgments

Z.G and R.J. was supported by the National Key Research and Development Program of China [2021YFF1200902] and [2023YFF1204802], and the National Natural Science Foundation of China [62203236 and 62273194]. Q.L., W.Z and W.H.W were supported by NIH grants R01 HG010359, P50 HG007735 and NSF DMS 1952386.

## Supplementary Materials

Text S1. Data splitting strategy for model training.

Text S2. System design and implementation of the web server. Text S3. Case application of the EpiGePT-online.

Text S4. Running time of the EpiGePT and baseline methods. Text S5. Implementation of Enformer model and Enformer+. Text S6. Data processing for ChromHMM annotation data.

Fig. S1. Three data partitioning strategies for model training and testing.

Fig. S2. Model architecture of EpiGePT for multiple epigenomic signals prediction. Fig. S3. EpiGePT’s performance in predicting DNase-seq and other epigenetic signals.

Fig. S4. Performance of EpiGePT and baseline methods on chromatin states classification, multiple epigenomic profiles prediction and causal variants classification.

Fig. S5. Ablation analysis of the EpiGePT model.

Fig. S6. Performance of EpiGePT in cross-cell-type prediction.

Fig. S7. The performance (auROC) of attention score of EpiGePT in distinguishing regulatory element-gene pairs at different distance ranges.

Fig. S8. Incorporating 3D genomic information from HiChip data enhances the predictive performance of EpiGePT on E-P regulatory interaction on K562 cell line.

Fig. S9. The fine-tuning performance of the EpiGePT model on predicting potential enhancer-promoter regulatory networks.

Fig. S10. The ROC and PR curves of the EpiGePT model on predicting potential enhancer-promoter regulatory networks.

Fig. S11. The GIS of ChIP-seq overlapped bins versus non-overlapped bins of POU5F1 centered at the TSS of ESRRB.

Fig. S12. Gene ontology enrichment analysis based on the top 5% TFs with high expression in ESCs.

Fig. S13. Case application of the EpiGePT-online.

Fig. S14. Enrichment result (Cellular component and Molecular function) of the nearest genes of the COVID-19 associated SNPs with the low LOS.

Table S1. The information of DNase-seq bam file across 129 biosamples from the ENCODE7 project.

Table S2. The information of RNA-seq tab-separated values (tsv) file across 129 biosamples from the ENCODE project.

Table S3. The information of DNase-seq, CTCF and other six Histone markers bam file across 28 cell lines or tissues from the ENCODE project (hg19).

Table S4. The information of DNase-seq, CTCF and other six Histone markers bam file across 104 cell lines or tissues from the ENCODE project (hg38).

Table S5. The information of RNA-seq tab-separated values (tsv) file across 28 cell lines or tissues from the ENCODE project (hg19).

Table S6. The information of RNA-seq tab-separated values (tsv) file across 104 cell lines or tissues from the ENCODE project (hg38).

Table S7. The preprocessed expression data of 711 human transcription factors from the ENCODE project across 129 biosamples.

Table S8. The preprocessed expression data of 711 human transcription factors from the ENCODE project across 28 cell lines or tissues (hg19).

Table S9. The preprocessed expression data of 711 human transcription factors from the ENCODE project across 104 cell lines or tissues (hg38).

Table S10. The order and names of epigenomes of the expression matrices across 56 epigenomes from the ROADMAP project.

Table S11. The preprocessed expression data of 642 human transcription factors across 56 epigenomes from the ROADMAP project.

