## Supplementary Texts S1-S6; Supplementary Figures S1-S14 for "EpiGePT: a Pretrained Transformer model for epigenomics"

* To whom correspondence should be addressed.

### Supplementary Texts

#### Text S1. Data splitting strategy for model training.

To comprehensively validate the performance of EpiGePT in predicting chromatin accessibility, we adopted three different data splitting strategies in the DNase[1] prediction experiment to verify the model's prediction ability when facing new genomic regions and cell types, which can meet researchers' usage needs to the maximum extent. Firstly, cross-cell type prediction refers to splitting the training and testing sets according to cell types in the same genomic region, where the cell types in the testing set have not appeared in the training set (Figs. S1b). Secondly, cross-genomic region prediction refers to splitting the training and testing sets according to genomic regions in the same cell type (Figs. S1a). Thirdly, simultaneous cross-cell type and genomic region prediction, where the prediction can be performed in completely novel cell types and genomic regions with the expression of transcription factors in that cell type. The training set needs to subset both cell types and genomic regions (Figs. S1c). To complete the latter two auxiliary predictions, we also split the data into 5 folds according to both cell types and genomic regions, so that both cross-validation can be performed in one round of training, but this will also reduce the amount of training and testing data.

#### Text S2. System design and implementation of the web server.

EpiGePT-online runs on a Linux-based Apache web server (https://www.apache.org) and utilizes the Bootstrap v3.3.7 framework (https://getbootstrap.com/docs/3.3/) for its web-frontend display. The backend of the server uses PHP v7.4.5 (http://www.php.net). The platform is compatible with the majority of mainstream web browsers, including Google Chrome, Firefox, Microsoft Edge, and Apple Safari.

#### Text S3. Case application of the EpiGePT-online.

The online prediction web service of EpiGePT enables users to predict eight types of epigenomic signals using EpiGePT without the need for setting up environments, writing code, or computational resources. In this section, we describe a usage scenario of EpiGePT-online for epigenomic signals prediction (Fig. S13). Users are provided with the flexibility to annotate either multiple genomic regions or a single locus at their discretion. Assuming an algorithmic researcher is interested in determining the potential regulatory role of a specific chromatin region based on its epigenetic modifications. In this case, the researcher can utilize EpiGePT-online to calculate the epigenetic signals on this region, to obtain references for assessing the potential regulatory role of the region. The submission prerequisites encompass two essential components. 1) The expression profiles of 711 TFs, which facilitate EpiGePT in acquiring precise cell type or tissue information. 2) The specific location of a locus on the genome or uploading of a bed file containing the information of genomic regions. It is worth noting that each line in the uploaded BED file should correspond to a 128kbp region to comply with the input length requirement of EpiGePT. If users select a specific locus, we will provide the predicted results for the region spanning 128kbp upstream and downstream of that locus. The web server allows users to upload expression values of 711 TFs in either numpy or comma-separated values (CSV) format. When predicting for $N$ genomic regions, users can obtain a downloadable matrix stored in CSV format with dimensions $(N \times 1000, 8)$. Each row denotes a 128bp genomic bin, and each column denotes an epigenetic profile. The specific referents of each row and column are provided in the downloadable table. This allows users to perform downstream analyses, such as related analyses in the areas of gene regulation and human disease.

### Text S4. Running time of the EpiGePT and baseline methods.

To demonstrate the computational efficiency of our model, we recorded the runtime of EpiGePT and baseline methods for one epoch on two sets of experiments, with different data sizes and input sequence lengths. Firstly, in the DNase signal prediction experiment on 129 cell types, with an input sequence length of 10kbp and using the same training data, Enformer requires approximately 3 hours and 4 minutes to complete one epoch, while EpiGePT only takes 2 hours and 17 minutes. In contrast, ChromDragoNN[2], which uses a genomic bin rather than a long region as the model input, requires 24 hours for pre-training and 8 hours for fine-tuning. In this case, the batch size of ChromDragoNN was set to 1024, which is equivalent to EpiGePT using a batch size of around 20. This modeling and computation approach presents challenges in terms of computational efficiency when dealing with large amounts of data. DeepCAGE[3] faces similar efficiency issues using the same approach. Even with a batch size of 256 on a single GPU, it still takes nearly 10 hours to complete one epoch of training. Secondly, we also recorded the running time of the models under larger-scale data and longer input sequences. When the number of input genomic bins increased from 50 to 1000, which corresponds to an input sequence length of approximately 128k, EpiGePT took approximately 3 hours to complete one epoch of training on 20 cell lines and 13,300 genomic regions, while Enformer required approximately 27 hours to train one epoch, as it required a longer input sequence of approximately 190kbp. Furthermore, EpiGePT without TF module (EpiGePT-seq) had approximately 1/4 of the parameters of Enformer and took approximately 2 hours and 40 minutes to train. In terms of performance, EpiGePT-seq performed similarly to Enformer on this dataset. This also explains why we chose to simplify the pure sequence model rather than directly adding a TF module to Enformer.

#### Text S5. Implementation of Enformer model and Enformer+.

To ensure a fair comparison between models and prevent the possibility of information leakage, we implemented the Enformer[4] model ourselves and trained it on our own collected data. Due to differences in dataset size and partitioning compared to Enformer, we reduced the number of encoder layers in Enformer to prevent overfitting. Thus, we reduced the number of encoder layers in Enformer to 3. Additionally, we introduced Enformer+ to enable a fair comparison between EpiGePT and Enformer in bin-level prediction. As Enformer takes only the DNA sequence as input, it tends to predict the same values for the same locus in different cell types, resulting in a loss of locus-level prediction ability. To address this, we incorporated the binding status and expression of the same transcription factors in Enformer+, and compared it to EpiGePT's performance on the same tasks.

### Text S6. Data processing for ChromHMM annotation data.

We downloaded the 15-state ChromHMM[5] annotations across 127 epigenomes from the ROADMAP project. The state of chromatin is annotated for each 200bp bin in a specific cell type. RNA-seq data of TFs across 56 cell types were download and extracted from the ROADMAP[6] project (Supplementary table S10 and S11). Subsequently, we mapped the 711 transcription factors to the downloaded RNA-seq data, resulting in the identification of RNA-seq data for 642 transcription factors. In the subsequent experiments, we utilized the expression data of these 642 transcription factors. We finally calculated the normalized TPM values of the 642 TFs on 56 cell types we extracted for the using in the classification model. For coarse grain chromatin state prediction, we took the state 'Quies' as low signal regions and other states as signal regions. For fine grain chromatin state prediction, we extracted the state 'TssA', 'TssAFlnk', 'TssBiv' and 'BivFlnk' as TSS regions, state 'EnhG', 'Enh' and 'EnhBiv' as enhancer regions, 'Quies' as low signal regions and other state as other regions. To balance the number of different chromatin states, we downsampled the low signal regions and obtained 921,074 bin each cell line finally.

### Supplementary Figures

#### Fig. S1

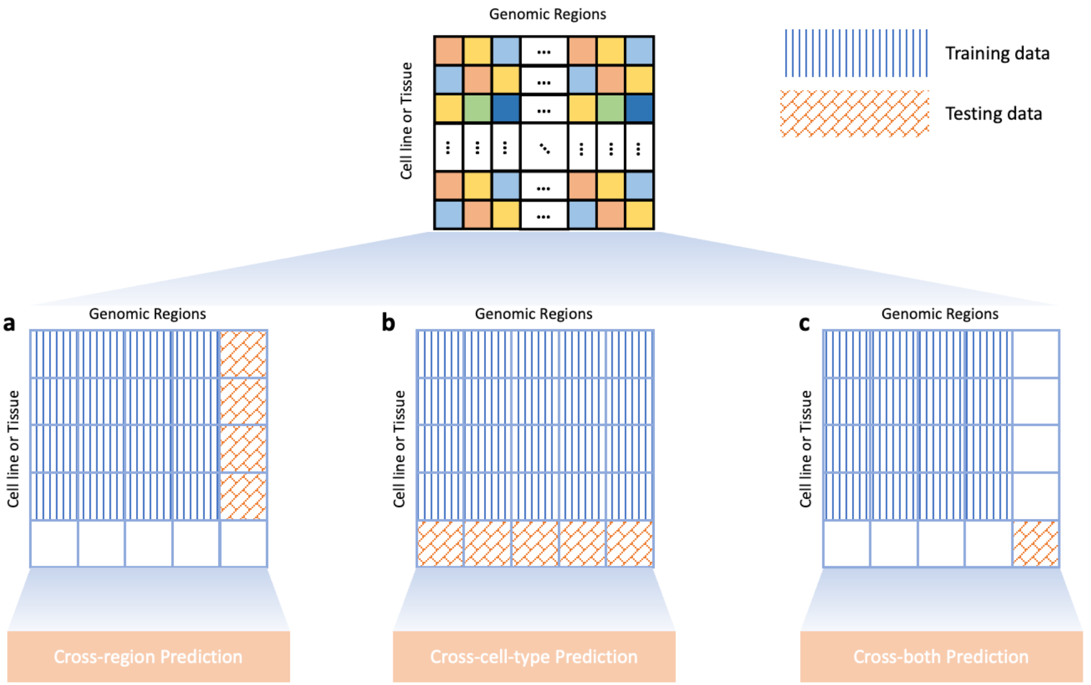

**Fig. S1. Three data partitioning strategies for model training and testing. a**, Cross genomic region prediction. The training and testing datasets utilized the expression profiles of identical cell types, but were evaluated on novel genomic regions for prediction. **b**, Cross cell type prediction. The training and testing datasets utilized the same genomic regions, but were evaluated on novel cell types for prediction. **c**, Cross genomic region and cell type prediction. The cell types and genomic regions used in the training and test sets were both different.

#### Fig. S2

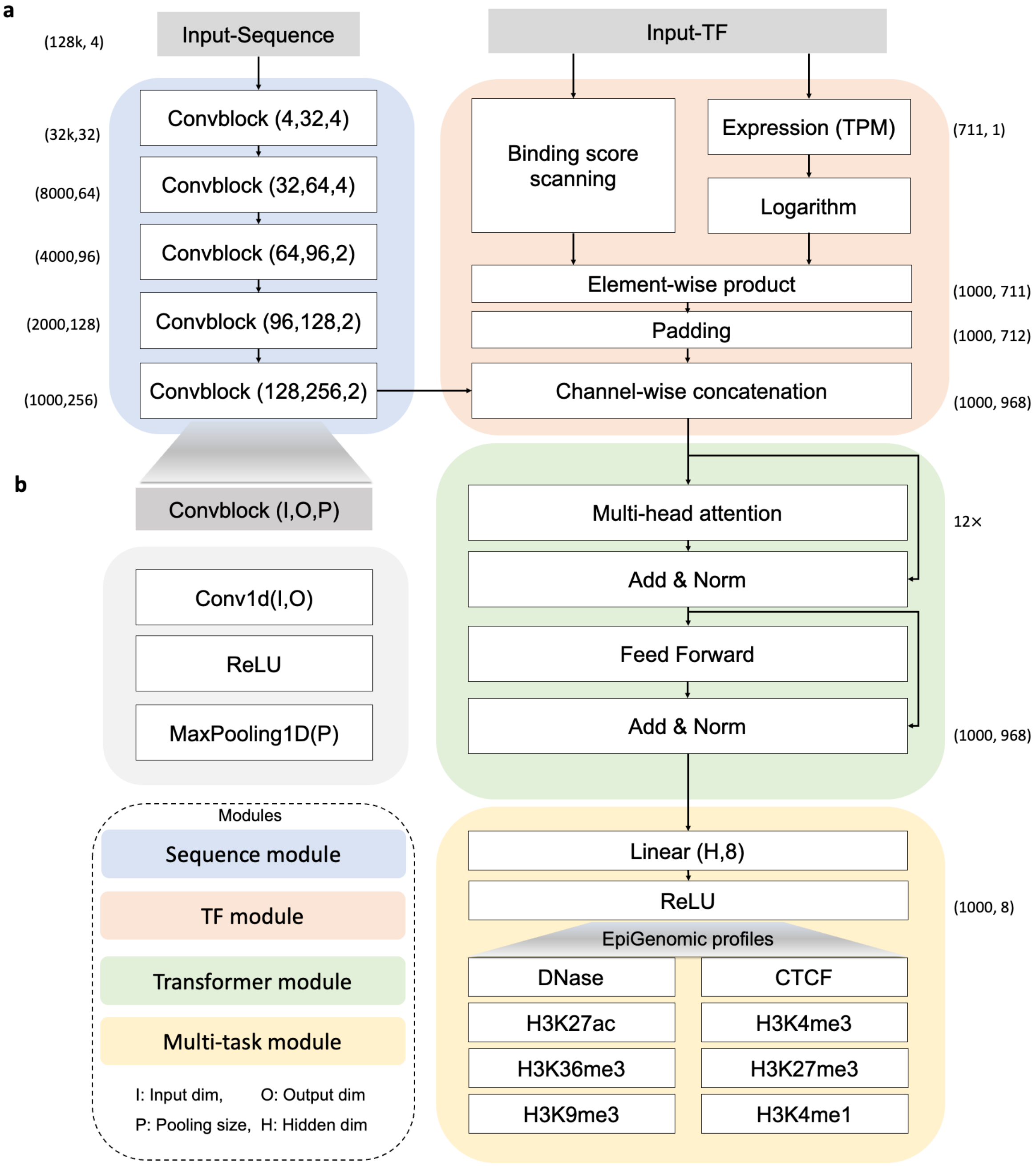

**Fig S2. Model architecture of EpiGePT for multiple epigenomic signals prediction. a**, The computational process of EpiGePT. The sequence module employs a stack of five convolutional layers followed by pooling operations, resulting in representations that capture sequence patterns. The TF module integrates motif binding information and gene expression data to represent cell-specific information. The Transformer module takes the genomic bin sequences mentioned above as input and learns the interaction relationships between bins, capturing the interactions among them. Finally, the obtained embeddings are mapped to the eight types of epigenomic signals through a fully connected layer. **b**, Specific details of the convolutional block involve the fusion of 1D convolution, ReLU activation function, and max pooling operation to achieve changes in the feature dimension O and extract bin-level features.

### Fig. S3

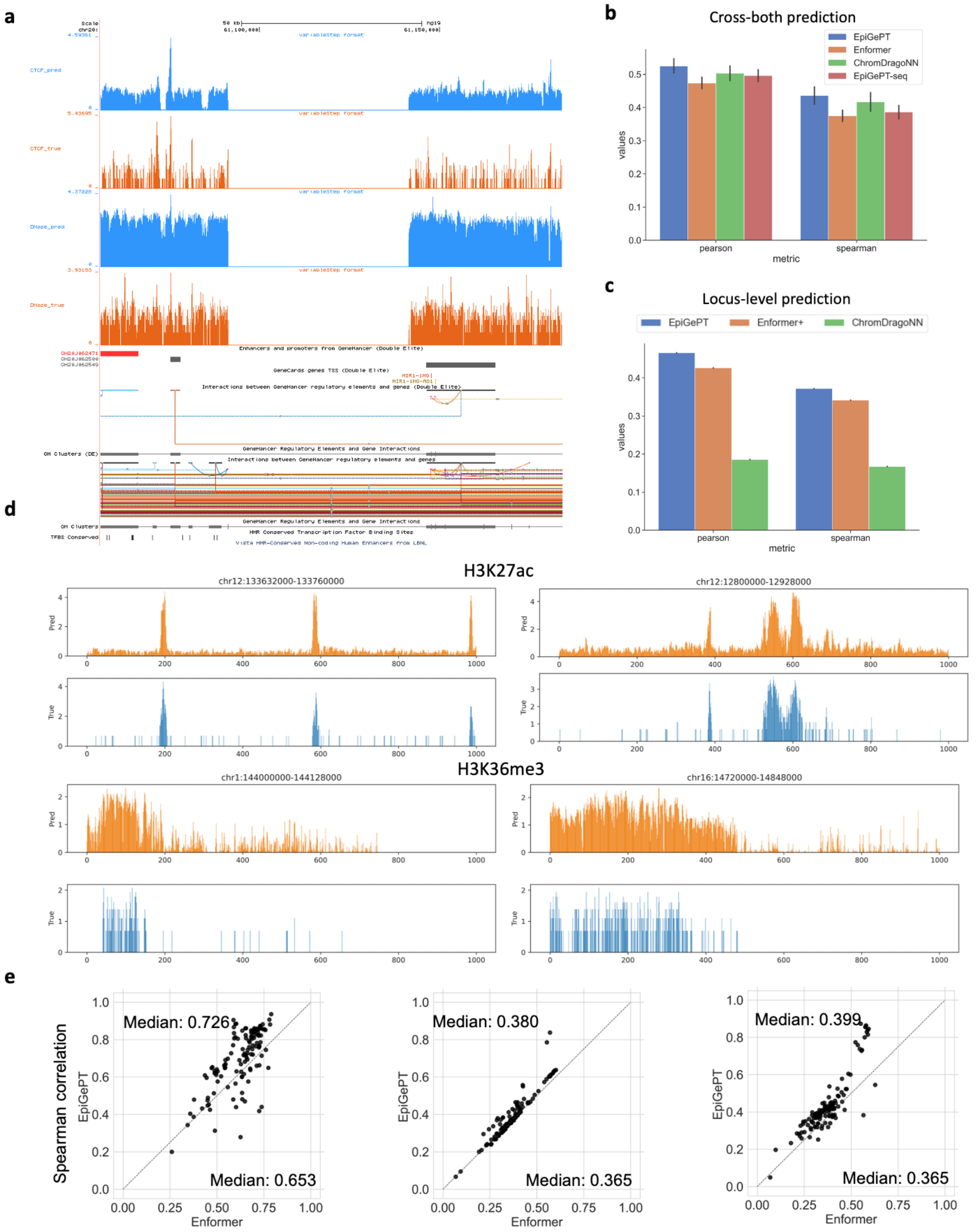

**Fig. S3. EpiGePT's performance in predicting DNase-seq and other epigenetic signals is demonstrated in a**, through visualization of predicted results for DNase and CTCF signals. EpiGePT is able to make accurate predictions for these signals, as well as for the regulatory relationships within a genomic region of 20th chromosome ranging from 61,100,000 to 61,150,000. **b**, EpiGePT and baseline methods were compared for their performance in predicting epigenetic signals in new cell types and genomic regions (cross-both prediction). The left panel shows the Pearson correlation coefficient, and the right panel shows the Spearman correlation coefficient. **c**, Locus level prediction of DNase signal. We predicted a value for each genomic locus, and calculated the correlation coefficient between the predicted values and true values for the same locus in different cell types. **d**, Visualization of predicted signals, such as the comparison between predicted and true values in a 128kbp region (from 133,632,000 to 133,760,000) on chromosome 12, shows that the presence of a large number of zeros in both the true and predicted signals can limit the correlation between the two signals. **e**, Comparison of EpiGePT and Enformer performance. Each point in the scatter plot represents the performance of Enformer on the data of a specific cell type (x-axis) compared to the performance of EpiGePT (y-axis). The three graphs represent the prediction of continuous DNase signals (spearman correlation coefficient).

#### Fig. S4

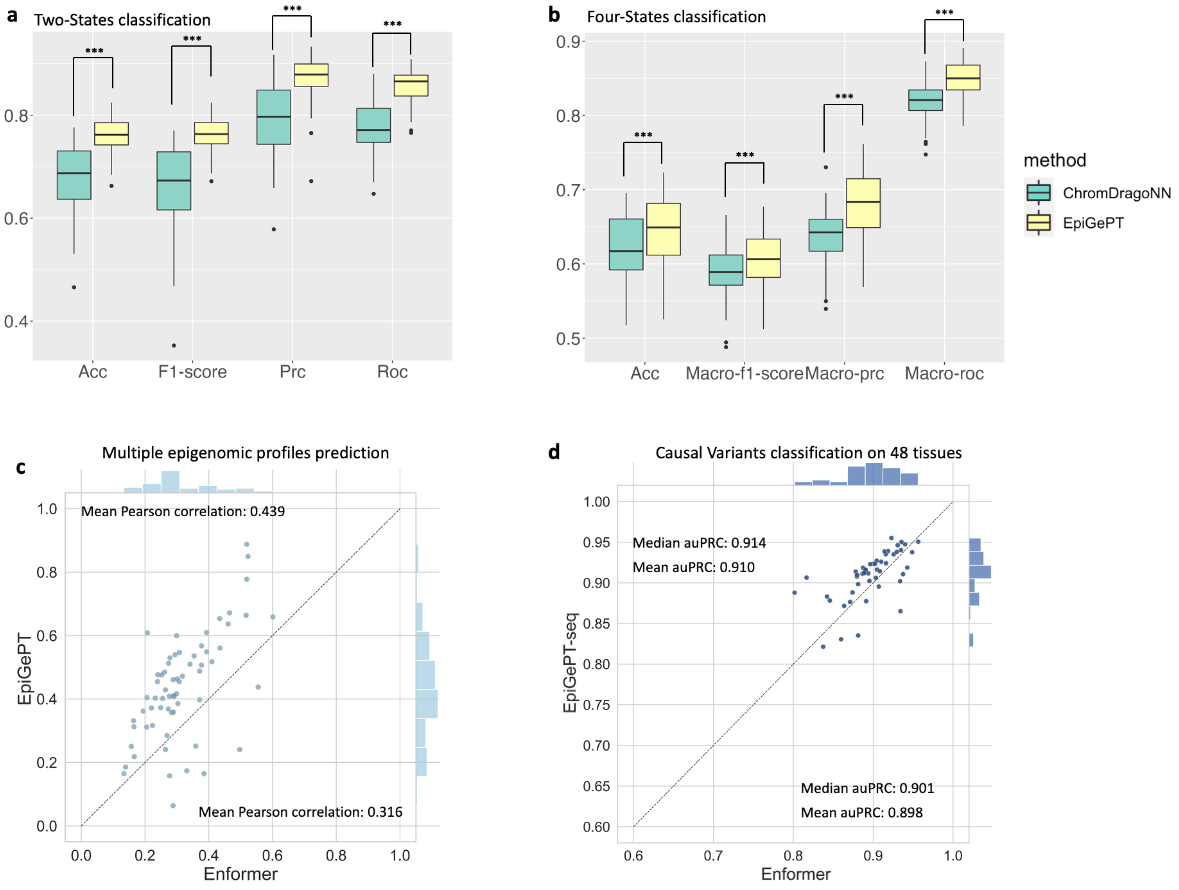

**Fig. S4. Performance of EpiGePT and baseline methods on chromatin states classification, multiple epigenomic profiles prediction and causal variants classification. a**, Binary classification of chromatin states for distinguishing functional regions on the chromatin based on the annotation data from ChromHMM-15-states. **b**, Four-class chromatin state classification is used to distinguish functional regions on the chromatin, including TSS, potential enhancers, other functional regions, and non-functional regions based on the annotation data from ChromHMM-15-states. *** indicates that the *p*-value is less than 1e-3 under one-sided Wilcoxon signed rank test. **c**, Cross-cell-type prediction of 8 epigenomic signals at 8 test cell types. Each dot denotes the Pearson correlation coefficient of the predicted signals and true signals at the specific cell types on a specific epigenomic signal. **d**, The performance of EpiGePT and Enformer in discriminating causal eQTLs across

48 tissues, each dot representing the average auPRC obtained from 5-fold cross-validation on a specific tissue.

#### Fig. S5

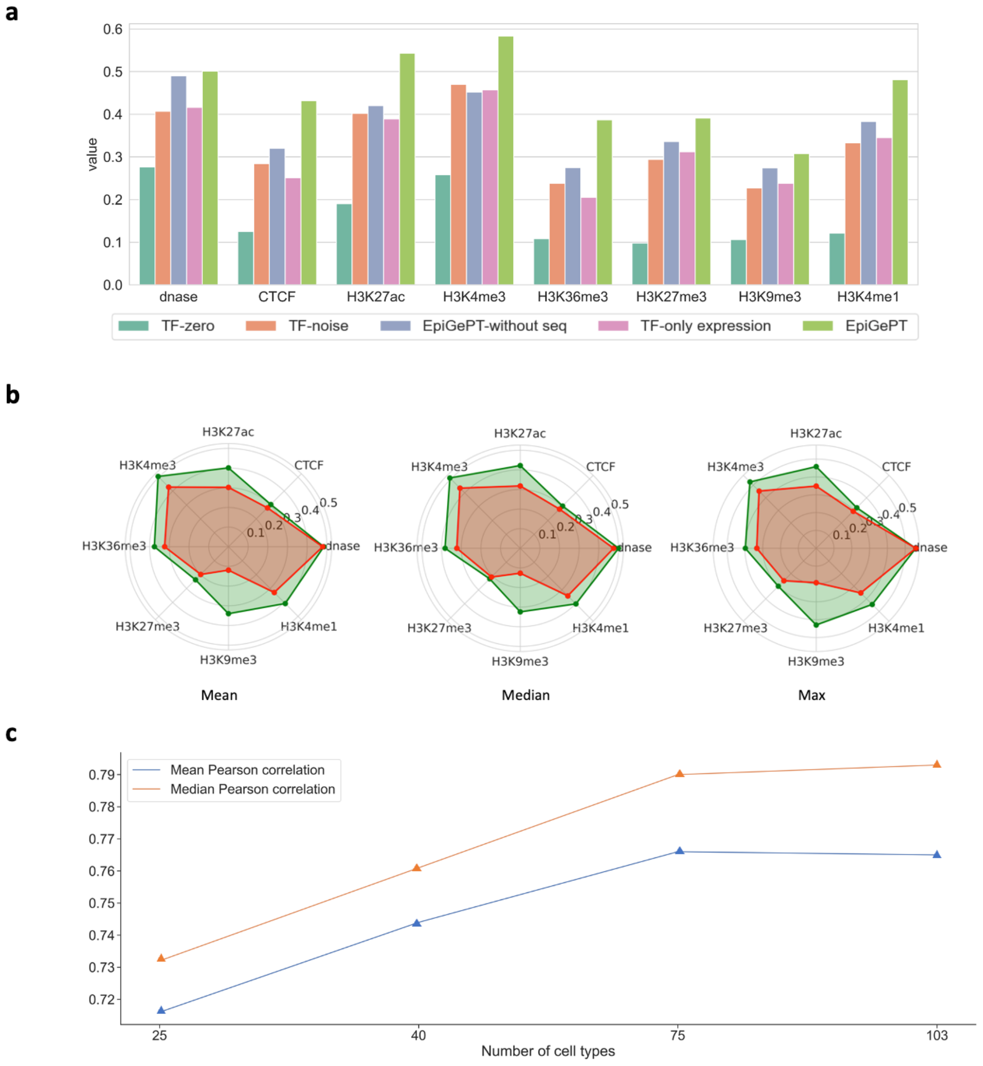

**Fig. S5. Ablation analysis of the EpiGePT model. a**, Ablation analysis on the TF module and the Sequence module, we observed a decrease in predictive performance for each module across eight chromatin epigenetic signals, as evidenced by a reduction in Pearson correlation coefficient. **b**, Ablation analysis on the Multi-task module. The green shaded area in the figure represents the results of multi-signal cross-cell-type predictions, while the red shaded area represents the results of training and predicting on each signal individually. It can be observed that the multi-task module has a positive effect on the model performance across all signals. **c**, Ablation analysis of the number of the training cell types. When the number of training cell types increases while the number of testing cell types remains constant, there is an increasing trend in performance as the number of training cell types increases.

#### Fig. S6

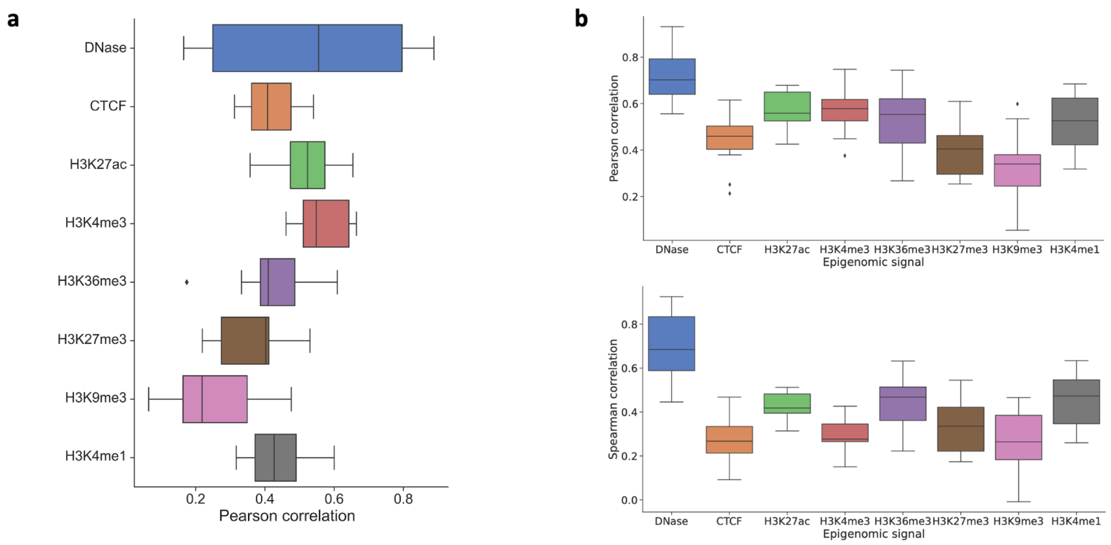

**Fig. S6. Performance of EpiGePT in cross-cell-type prediction. a**, The predictive performance of EpiGePT on 8 unseen cell types on hg19 reference genome (pearson correlation coefficients). **b**, The predictive performance of EpiGePT on 19 new cell types on hg38 reference genome (upper: pearson correlation coefficients, lower: Spearman correlation coefficients).

#### Fig. S7

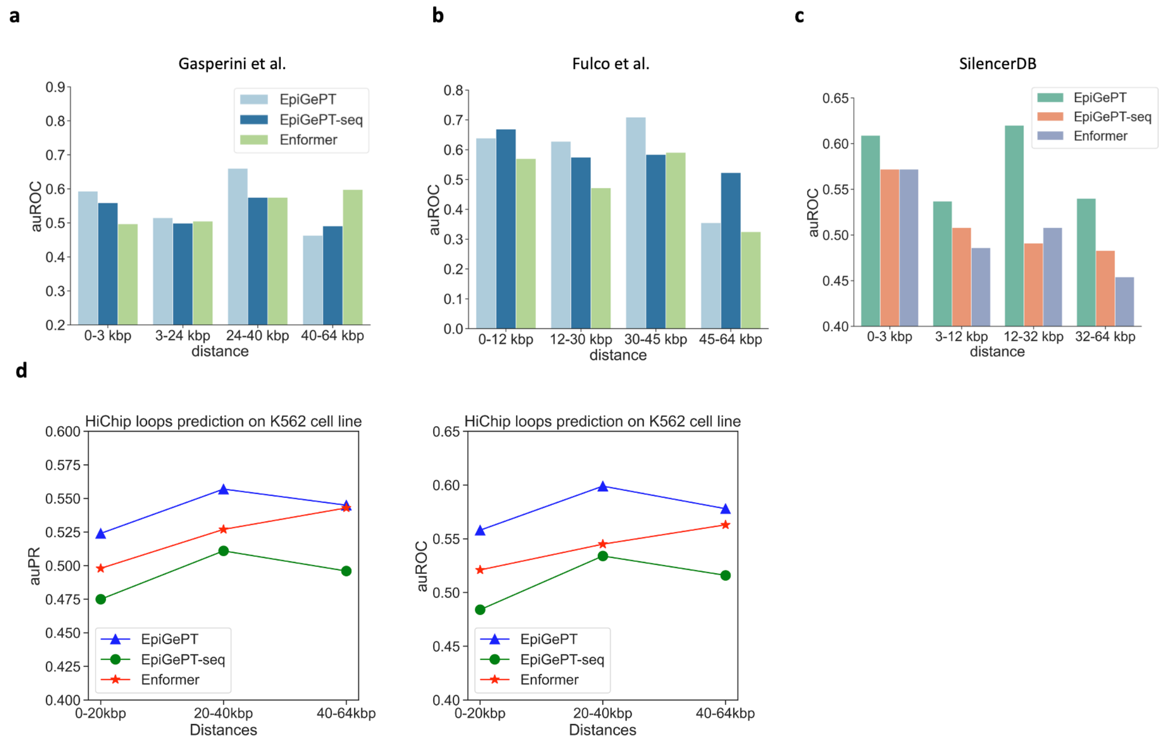

**Fig. S7. The performance (auROC) of attention score of EpiGePT in distinguishing regulatory element-gene pairs at different distance ranges. a**, The performance of EpiGePT in distinguishing enhancer-gene pairs at different distance ranges on the data from Gasperini et al[7]. **b**, The performance of EpiGePT in distinguishing enhancer-gene pairs at different distance ranges on the data from Fulco et al[8]. **c**, The performance of EpiGePT in distinguishing silencer-promoter pairs at different distance ranges on the data from SilencerDB[9]. **d**, The performance (auROC and auPR) of attention score of EpiGePT in distinguishing HiChIP loops of H3K27ac at different distance ranges on K562 cell line.

#### Fig. S8

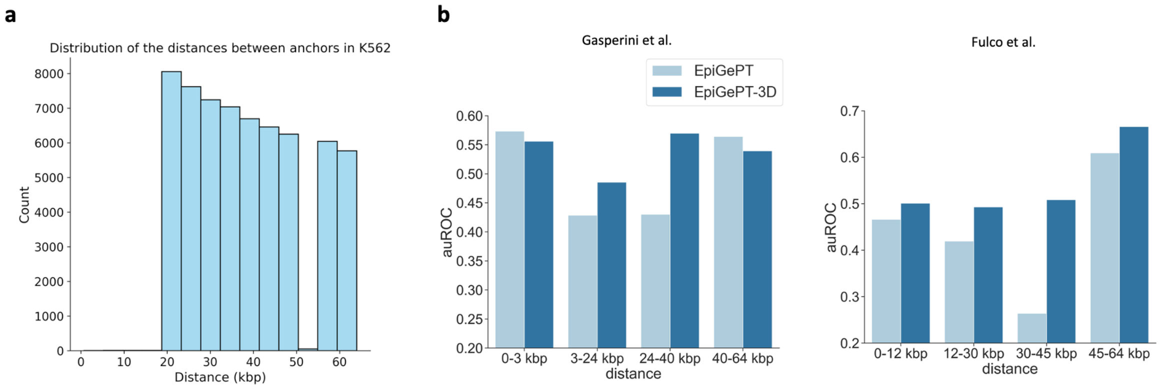

**Fig. S8. Incorporating 3D genomic information from HiChip data enhances the predictive performance of EpiGePT on E-P regulatory interaction on K562 cell line. a**, The distance distribution between the two anchors of the filtered loops on the K562 cell line. **b**, The performance (auROC) of self-attention scores of EpiGePT and EpiGePT-3D in identifying enhancer-promoter interactions across different distance ranges on the K562 cell type.

### Fig. S9

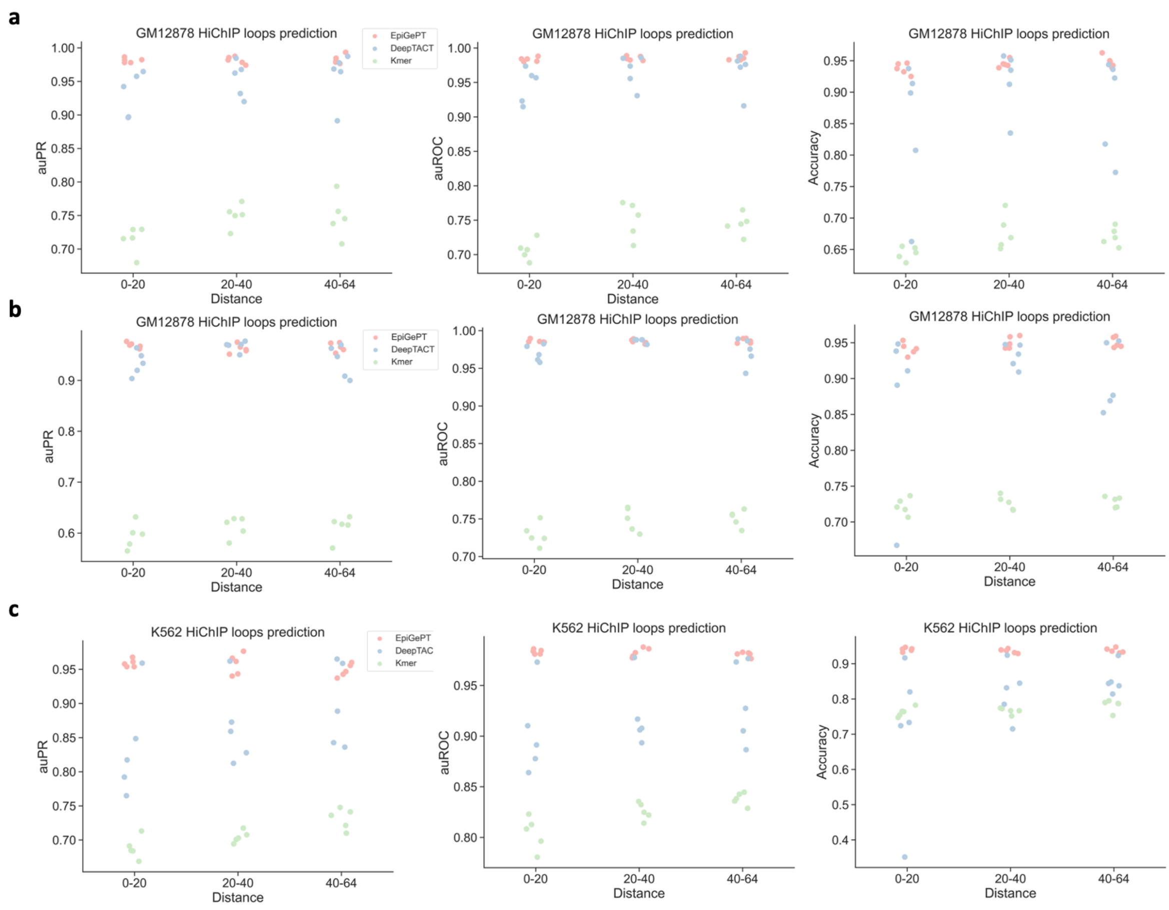

**Fig. S9. The fine-tuning performance of the EpiGePT model on predicting potential enhancer-promoter regulatory networks. a**, The performance (measured by auROC and auPRC) of the fine-tuned EpiGePT model and baseline methods (DeepTACT and Kmer) on HiChIP loops data in distinguishing enhancer-gene pairs at various distance ranges (0-20 kbp, 20-40 kbp and 40-64 kbp). **b**, The performance of the fine-tuned EpiGePT model and baseline methods on HiChIP loops data in distinguishing enhancer-gene pairs under 1:2 positive-negative sample ratio on GM12878 cell line. **c**, The performance of the fine-tuned EpiGePT model and baseline methods on HiChIP loops data in distinguishing enhancer-gene pairs under 1:2 positive-negative sample ratio on K562 cell line.

#### Fig. S10

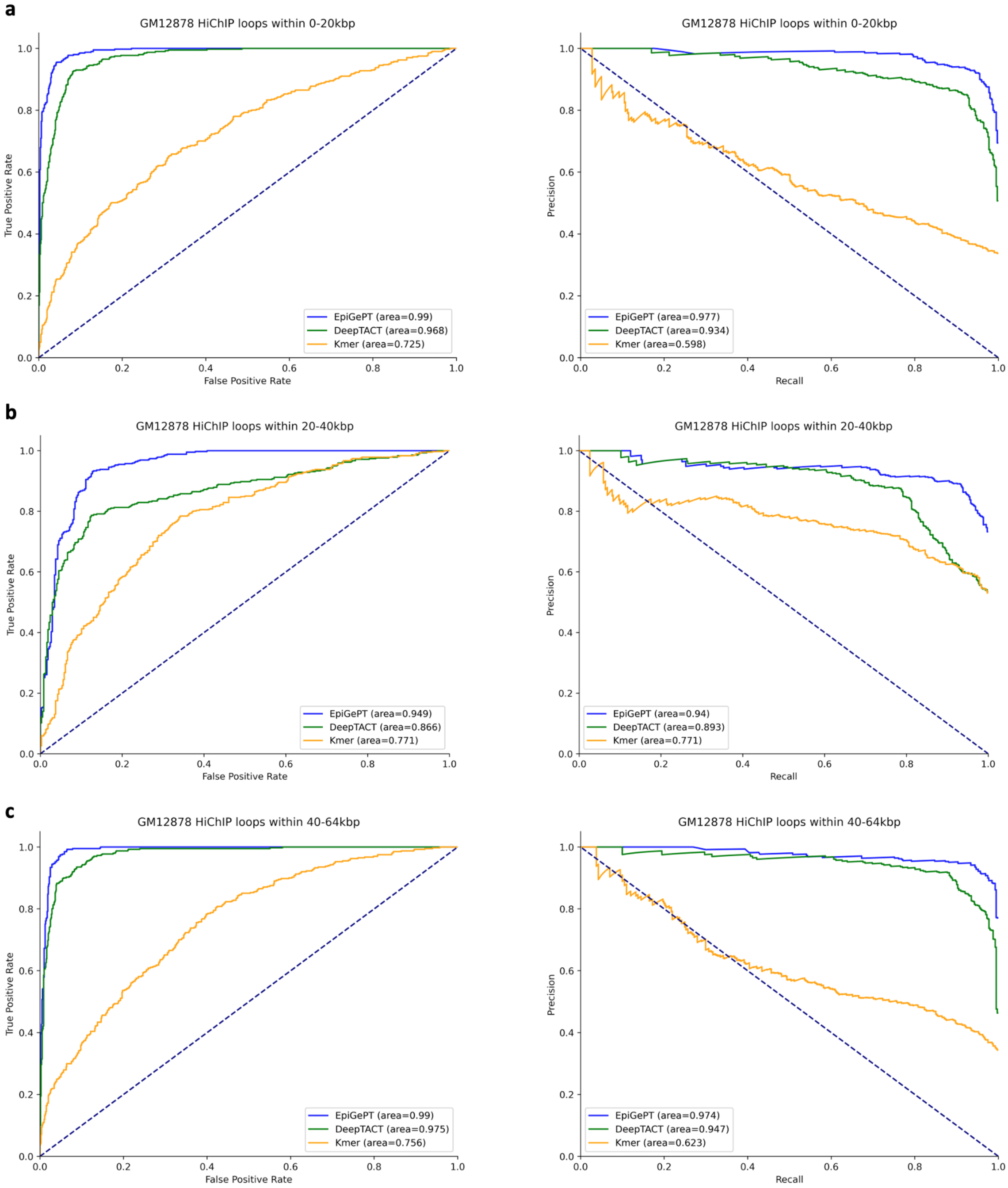

**Fig. S10. The ROC and PR curves of the EpiGePT model on predicting potential enhancer-promoter regulatory networks. a**, The ROC and PR curves of EpiGePT and baseline methods for predicting HiChIP loops from the GM12878 cell line (0-20 kbp). **b**, The ROC and PR curves of EpiGePT and baseline methods for predicting HiChIP loops from the

GM12878 cell line (20-40 kbp). **c**, The ROC and PR curves of EpiGePT and baseline methods for predicting HiChIP loops from the GM12878 cell line (40-64 kbp).

### Fig. S11

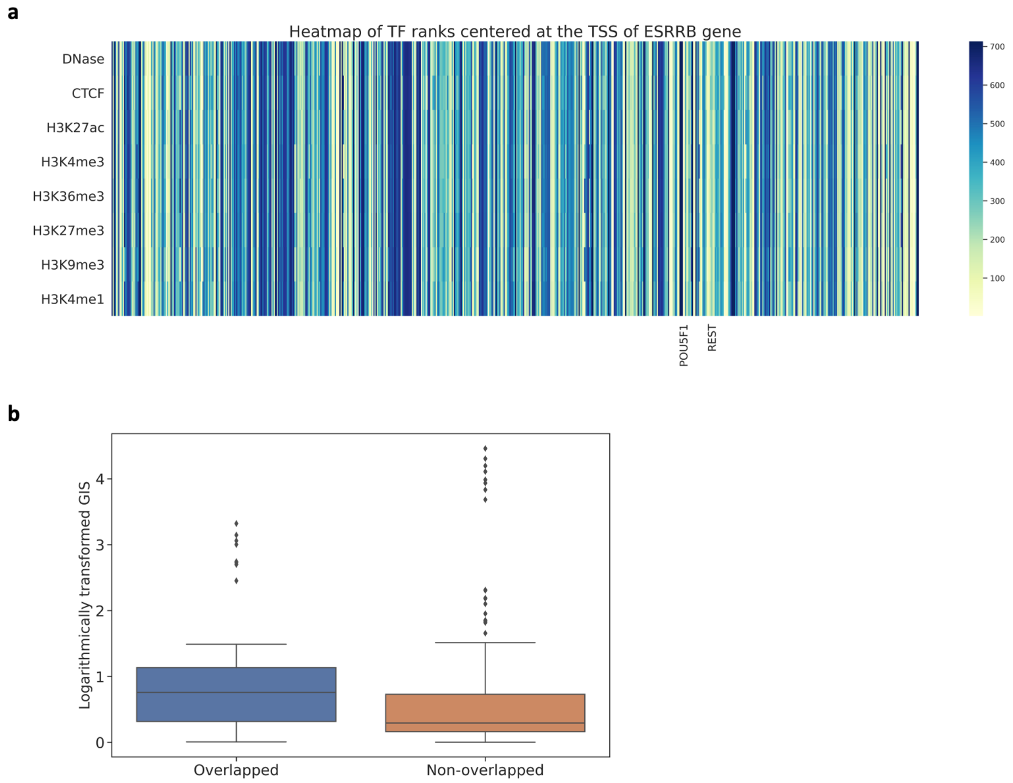

**Fig. S11. The GIS of ChIP-seq overlapped bins versus non-overlapped bins of *POU5F1* centered at the TSS of *ESRRB*. a**, Heatmap of TF ranks across 128 kbp region surrounding the TSS of *ESRRB* gene, each row denotes an epigenomic signal and each column denotes a TF. **b**, Distribution of non-zero GIS values on overlapped and non-overlapped bins in chip-seq data (ENCFF696NWL).

### Fig. S12

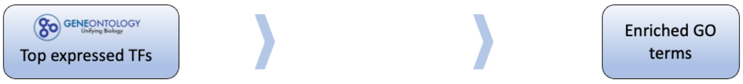

| GO Term ID | Biological process | P-value | FDR |
|---|---|---|---|
| GO:0048598 | embryonic morphogenesis | 8.78e-07 | 1.98e-04 |
| GO:0009790 | embryo development | 1.92e-07 | 4.99e-05 |
| GO:0048568 | embryonic organ development | 1.06e-07 | 2.86e-05 |
| GO:0009792 | embryo development ending in birth or egg hatching | 1.95e-04 | 1.79e-02 |
| GO:0030154 | cell differentiation | 6.67e-07 | 1.58e-04 |
| GO:0001892 | embryonic placenta development | 2.09e-05 | 2.92e-03 |

**Fig. S12. Gene ontology enrichment analysis based on the top 5% TFs with high expression in ESCs.** The results showed lower significance for biological processes associated with embryonic cell development compared with GO terms enriched with the top 5% ranked TFs.

#### Fig. S13

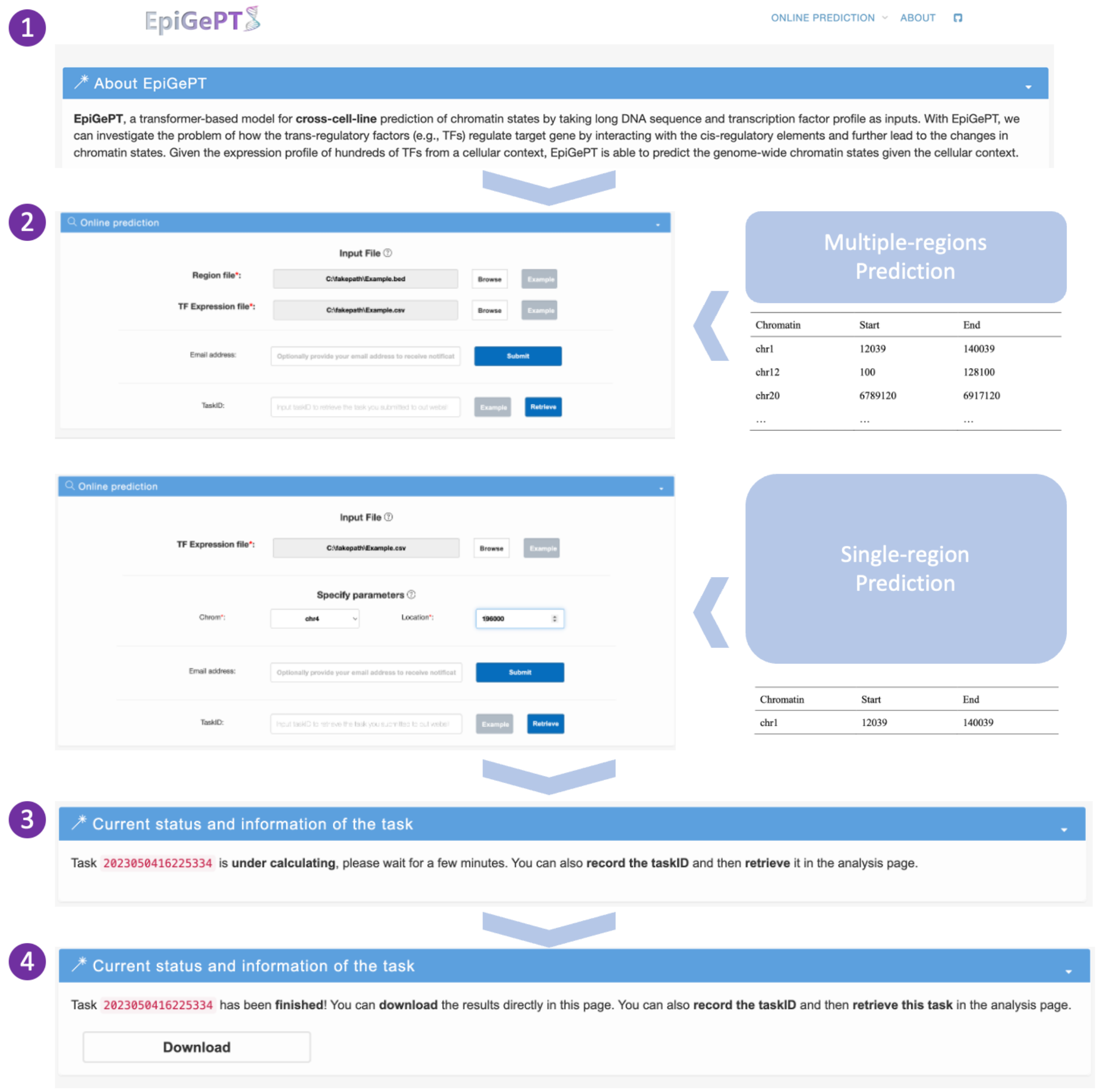

**Fig. S13. Case application of the EpiGePT-online.** Users can choose either single locus annotation or multi-region annotation on EpiGePT-online, and each genomic region requires a length of 128kbp. Users need to upload the TPM values of transcription factors expression simultaneously. After annotation, users can enter the result page and download the predicted files. The predictions are provided at the resolution of 128bp genomic bins, and users can obtain the predicted signals for these eight epigenomic profiles. Additionally, users have the option to download the prediction results in CSV format for further analysis and exploration.

### Fig. S14

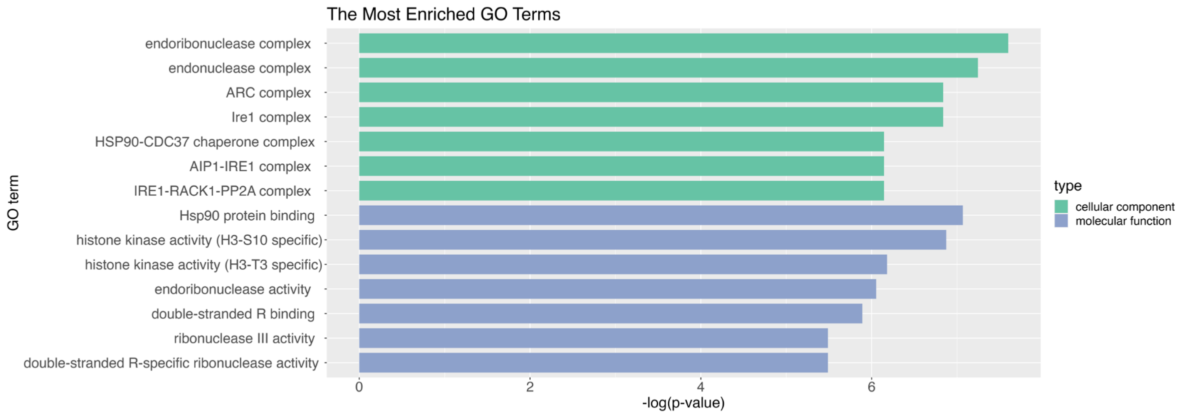

**Fig. S14 Enrichment result (Cellular component and Molecular function) of the nearest genes of the COVID-19 associated SNPs with the low LOS.**

#### Supplementary Tables

Table S1. The information of DNase-seq bam file across 129 biosamples from the ENCODE[10] project.

Table S2. The information of RNA-seq tab-separated values (tsv) file across 129 biosamples from the ENCODE[10] project.

Table S3. The information of DNase-seq, CTCF and other six Histone markers bam file across 28 cell lines or tissues from the ENCODE[10] project (hg19).

Table S4. The information of DNase-seq, CTCF and other six Histone markers bam file across 105 cell lines or tissues from the ENCODE[10] project (hg38).

Table S5. The information of RNA-seq tab-separated values (tsv) file across 28 cell lines or tissues from the ENCODE[10] project (hg19).

Table S6. The information of RNA-seq tab-separated values (tsv) file across 105 cell lines or tissues from the ENCODE[10] project (hg38).

Table S7. The preprocessed expression data of 711 human transcription factors from the ENCODE[10] project across 129 biosamples.

Table S8. The preprocessed expression data of 711 human transcription factors from the ENCODE[10] project across 28 cell lines or tissues (hg19).

Table S9. The preprocessed expression data of 711 human transcription factors from the ENCODE[10] project across 105 cell lines or tissues (hg38).

Table S10. The order and names of epigenomes of the expression matrices across 56 epigenomes from the ROADMAP[6] project.

Table S11. The preprocessed expression data of 642 human transcription factors across 56 epigenomes from the ROADMAP[6] project.
